# Using rotational integration of oblique interferometric scattering (RO-iSCAT) to track axial spatiotemporal responses of membrane protrusions

**DOI:** 10.1101/2025.03.23.644841

**Authors:** Junyu Liu, Yean Jin Lim, David Herrmann, Paul Timpson, Tri G. Phan, Huafeng Liu, Min Guo, Woei Ming Lee

## Abstract

Despite the crucial importance of dynamic membrane protrusions for understanding phagocytosis, cellular communication and mechanobiology, current imaging modalities struggle to quantitatively track their real-time, 3D spatiotemporal dynamics with sufficient molecular specificity and minimal perturbation. Many membrane protrusions studies still utilize confocal microscopy where its axial resolution and high phototoxicity remains a key limiting factor for live axial imaging. We discovered that multiple rotational oblique interference scattering (RO-iSCAT) leverages off-axis illumination to induce a larger lateral shift in out-of-focus iSCAT signals compared to in-focus signals. This phenomenon provides a foundation to generate speckle-free widefield interferometric signals with a 10-fold signal to noise ratio improvement, eliminating the need for any background subtraction. RO-iSCAT enables real-time, label-free, and minimally invasive imaging of diverse membrane protrusions within complex co-cultures. RO-iSCAT enables nanoscale-sensitive tracking of membrane protrusion dynamics along the axial direction. This allows for the construction of dynamic axial variance maps, facilitating quantitative measurements of membrane protrusion formation at tens to hundreds of nanometer displacements, without requiring 3D volumetric imaging. RO-iSCAT empowers real time quantitatively dissection of the axial spatiotemporal complexities of membrane protrusions and unlock future insights into fundamental processes like cell migration, durotaxis, and intercellular communication.

**Key points:**

- Discovered that multiple integrated rotational oblique interference scattering (RO-iSCAT) generates speckle-free widefield interferometric signals with a 10-fold signal to noise ratio improvement, eliminating the need for any background subtraction.
- Removed need for 3D volumetric imaging to quantified axial motion of membrane protrusion forming tethers, trails and bridge with within ∼ tens of nanometer accuracy.
- Enabled classification of membrane protrusions that, despite possessing identical chemical compositions, are differentiated by their interactions, thus offering a qualitative comparison of membrane protrusions at the nanoscale in living cells.

**Summary Figure:** 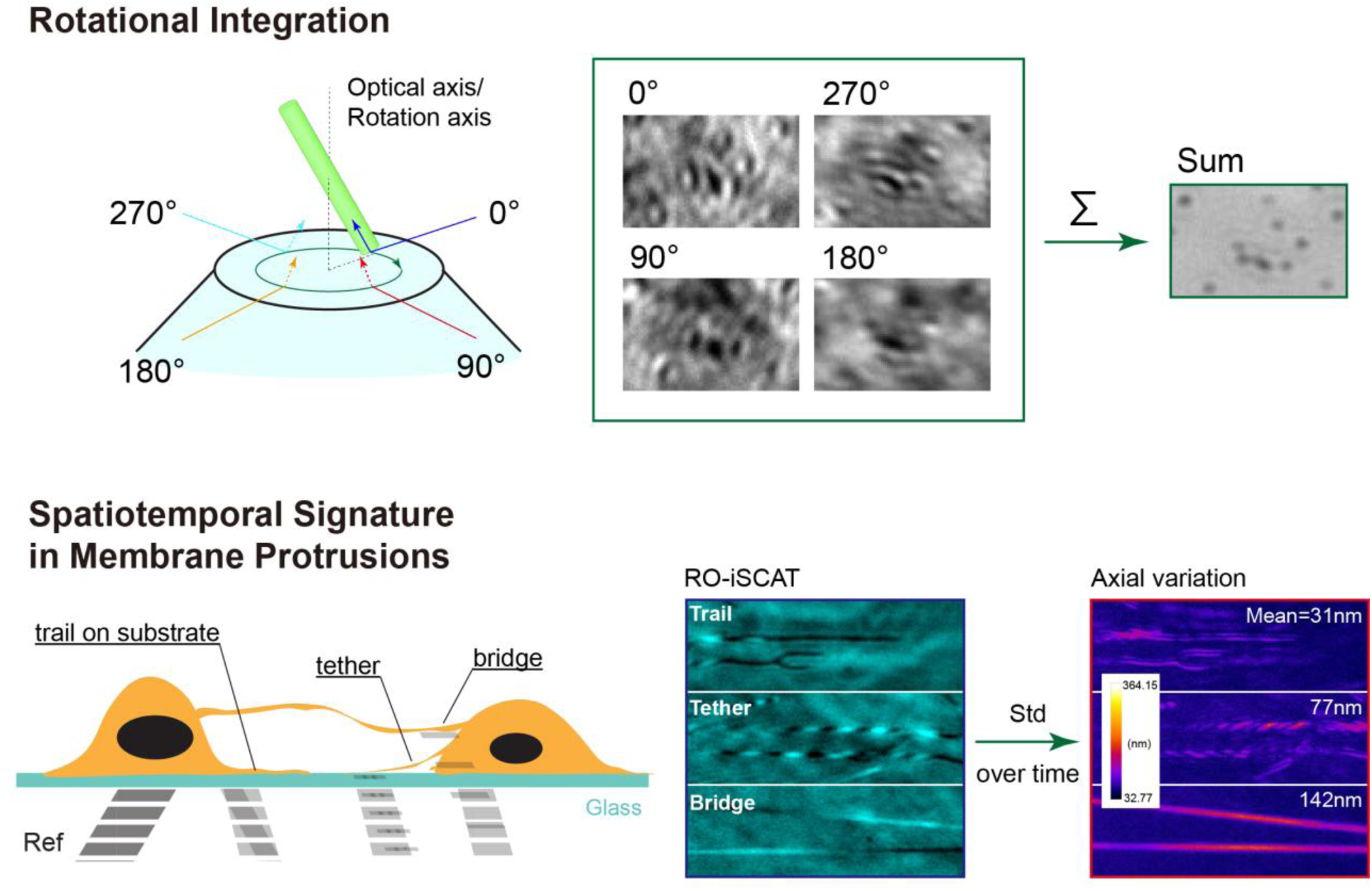

## Introduction

Membrane protrusions (lamellipodia, pseudopodia, filopodia, microvilli, invadopodia, and podosomes) possess dynamic three-dimensional spatiotemporal behaviours because they mediate a wide variety of extracellular interactions between a cell and its three-dimensional microenvironment ^1^. While these dynamic protrusions are the result of cytoskeletal (e.g. actin, microtubules) rearrangement, their 3D spatiotemporal relationship are initiated by the activation and clustering of membrane receptors ^2^. In particular, 2D spatiotemporal tracking of filopodia- membrane extensions indicate their role in mechanical and chemical sensing ^3^, phagocytosis ^4, 5^ and migration ^6, 7^. Observations of protrusion dynamics on coated substrates, between cells and in tissue ^8^ have led to discoveries on contact-dependent cell-cell communication ^9^, twisted tethers ^10^, forming migrasomes from retraction fibers ^11^, and tunnelling nanotube ^12^ as well as gaining closer insight into tissue development ^13^.

Existing tools to track membrane protrusion, extensions and distribution in the spatiotemporal domain relies heavily on standard light microscopy technique (brightfield-phase contrast, fluorescence), have inadequate resolution, are prone to phototoxicity, and lacking specific fluorescence markers, cannot readily classify the transient behaviors of 3D membrane protrusion and extensions in live cell cultures. Whilst the use of volumetric imaging technologies ^14^ and advanced image processing ^15^ has made significant advances, the issue of phototoxicity and photobleaching remains a concern for longitudinal imaging ^16^ that is necessary for quantitative mapping of membrane protrusions. Electron microscopy (EM), on the other hand, has become a routine tool to identify these protrusions that forms membrane bridges because of its ability to measure physical feature membrane protrusion based on physical size, diameter (50–200 nm diameter), the distance between distant cell and importantly, and their proximity with substrate for classification ^17^. Unfortunately, EM slices face methodological difficulties because membrane protrusion such as tethers and tunnelling nanotubes are often fragile after chemical fixation, and prone to deform due to sample preparation. Owing to well-defined refractive index difference in actin and lipid in thin membrane protrusion, it is plausible that interference scattering microscopy (iSCAT) ^18–20^ signals can be more effectively at quantitative tracking the transient movements of different types of membrane protrusions that form, disassemble and maintain at the nanoscale in 3D than fluorescence microscopy. However, iSCAT signals often require background subtraction ^20^, which can be challenging to implement and may fail to remove speckles in populated cell cultures, thus hindering the tracking of protrusion spatiotemporal patterns in live cells.

This paper examines the optical principles of a Rotational Oblique Interferometric Scattering signals (RO-iSCAT) to achieve speckle free interferometric scattering signals in real time. We then follow on to explain how RO-iSCAT interference patterns are used to track spatiotemporal of membrane protrusion; that can transits into trails (i.e. retraction fibers to migrasomes), membrane tethers or bridges ^21^. We provided evidence demonstrating that axial variation maps of interference scattering signals are effective in accurately categorizing various membrane protrusions. The axial variation information possesses rich spatial temporal signature even within a single protrusion that surpass standard kymograph in fluorescence or scattering images which only consider rudimentary shapes. The paper comprises of 4 main parts – numerical model of RO-iSCAT, quantification of imaging resolution of RO-iSCAT, identification of membrane protrusion using spatial signature in lateral interference pattern alongside with variance in axial displacement analysis of RO-iSCAT images, and longitudinal tracking of membrane protrusions in co-cultured cells lines.

## Results

### Rotational integration removes out-of-focus interference scattering signals

Under off-axis illumination, we observed that out-of-focus iSCAT signal experienced a larger lateral shift than in-focus iSCAT signal (**Supplementary Video 1**). To explain this effect, we began with a numerical model (**Supplementary Note 1** and **Methods**) and synthesized the interference intensity signals of RO-iSCAT. The scheme of oblique illumination we adopted was at a single angle, where an incoming illumination (blue line and shades) entered the sample at an oblique angle 𝜃 along a single azimuthal orientation 𝜑 (**Fig. 1a**). The path length difference arises from refractive index difference between glass-water interface (reference field) and scattered light that is necessary to form an iSCAT signal. Whilst the reference field is constant (**Fig. 1a**, first reflecting surface-blue line), the scattered signal (**Fig. 1a**, green line) varies along the axial plane 𝑧. Because iSCAT signal is of interferometric nature, the signal is changed by the properties of scattered signal collected by the imaging lens. This phase delay (defocused wavefront) changes with axial distance 𝑧 of the imaging lens for each oblique angles 𝜃 and azimuthal direction 𝜑.

**Fig. 1.**
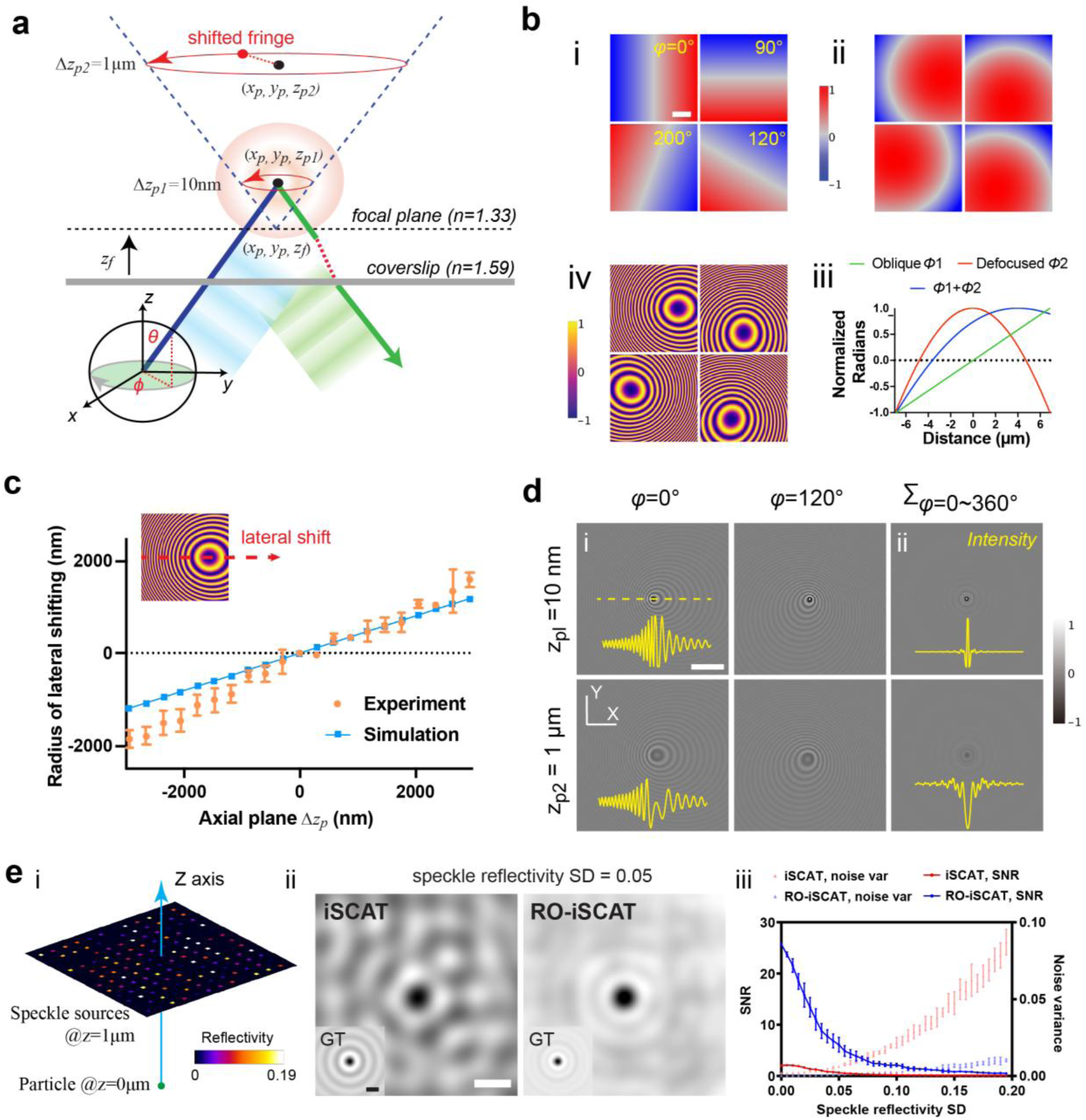
Modelling and simulation of RO-iSCAT. **a)** Schematic diagram of incident, reflection, scattering fields, and several reference planes in azimuthal iSCAT. **b)** Numerical simulation with 14 microns FOV and 70 nm step based on the modelling. Including i, off-axis oblique phase and ii, oblique convolved with defocused phase, respectively at 0°/90°/120°/200° with a 10 μm defocused length and 22° oblique angle. iii, Profiles of off-axis oblique, defocused, and total phase difference along the horizontal central axis of FOV. iv, Interference pattern at corresponding azimuth angle. **c)** The radius of lateral shifting under defocused length ranging from -2.5 μm to 2.5 μm in ∼290 nm step extracted from simulation and experimental data. The experiments were repeated five times independently, as indicated by error bars (mean +/- SD). **d)** i, Circumferential lateral-shifting related to illumination azimuth and defocused length. ii, Azimuthally integrated interference pattern. Also attached the corresponding intensity profile along the horizontal central axis of FOV. **e)** i, Sketch presentation to show the strategy of simulating different speckle noise level. One single particle at in-focus plane as signal source, while a series of speckles are placed at 10-micron depth with their reflectivity value following the Gaussian distribution. ii, In-focus fringes from iSCAT versus RO-iSCAT based on the object in i. Inserted ground truths are pure interference fringe only from the in-focus particle. iii, Noise variance and fringe SNR curve with varying reflectivity levels of speckle sources, both in iSCAT versus RO-iSCAT. **Scale bars**: **b), d)** 3 μm, **e)** 1 μm.

At each oblique angle, an off-axis phase shift causes iSCAT interference fringe pattern to shift laterally away from the focal plane. We numerically calculated the phase and fringe shifts at different azimuthal angle (**Fig. 1b**, **Supplementary** Fig. 1 and **Supplementary Video 2**). Each off-axis oblique phase delay (**Fig. 1b** i) from different azimuthal sources (𝜑 = 0°, 90°, 120°, 200°) is convolved with defocusing phase delay that will create nonlinear phase shifts (**Fig. 1b** ii, iii), which directly translate to the intensity fringes translating laterally (**Fig. 1b** iv).

To confirm the effect of lateral shifts in intensity fringes in RO-iSCAT, first we simulated lateral shift of intensity fringes across multiple axial planes from -2 µm to 2 µm and compared with the measured experimental results (**Supplementary Video 3** left). From both our model and experiment (**Fig. 1c**), we observed that the lateral fringe shifts increased further away from the focal plane, whereas at the focal plane, the iSCAT signal experience shift almost negligible. Then, we examined the modelled shift in fringes along the transvers plane at azimuthal rotational direction 𝜑 = 0° and 120° (**Fig. 1d** i) as well as the final integrated RO-iSCAT images 𝜑 = 0° − 360° (**Fig. 1d** ii). The integrated RO-iSCAT shows a significant reduction of side lobe (profiles in **Fig. 1d**) that indicates an increased visibility of the interference fringes at the focal plane. This rotational oblique configuration reduces out-of-focus signal equivalent to confocal configuration^19^ (**Supplementary** Fig. 2 and **Supplementary Video 4**). Hence, directly integrating multiple oblique illuminated iSCAT signals generates a high contrast iSCAT image at the focal point only, without intensity losses that can occur in background subtraction ^20^ or pinholes ^19^.

Next, we turned to examine the 3D interference point spread function (iPSF) of on axis (𝜃 = 0.5°), iSCAT and off axis ( 𝜃 = 22°), RO-iSCAT over a 4×4×8-micron FOV under 1.4 NA objective lens (**Supplementary** Fig. 3a) through a Boundary-Element-Method platform ^22^. RO- iSCAT possess a higher lateral signal to noise ratio and narrower expansion than conventional iSCAT (**Supplementary** Fig. 3b and 3c). To quantity the improvement on noise rejection and SNR, we synthetically generated fringe by 1) the pure signal from a single particle at focal plane alongside with 2) different levels of speckle noise from an out-of-focus plane (**Fig. 1e** i and **Methods**). RO-iSCAT effectively rejected most of the speckle artifacts from background (**Fig. 1e** ii) thus improved the SNR from 0.49 (**Fig. 1e** ii left) significantly to 5.65 (**Fig. 1e** ii right). Even under increasing noise conditions, RO-iSCAT consistently achieve high SNR when compared to background substraction (**Fig. 1e** iii, light blue versus light red scatters) and a ten-fold higher image SNR (Fig. 1e iii, blue versus red curve. **Supplementary** Fig. 4).

### RO-iSCAT imaging achieve speckle-free iSCAT without needing background subtraction

The role of background subtraction in majority of iSCAT methods ^19, 20, 23–26^ is to remove all extrinsic factors (i.e. non-uniform illumination, and unwanted coherent noise and interference) so as to increase SNR and reach single protein sensitivity. Background subtraction uses iSCAT images recorded with no focal drift and ultra clean glass slides that is free of any sample feature ^20^. In this section, we compared SNR of RO-iSCAT versus standard background subtraction (iSCAT). The sample was a glass coverslip dish containing surface-bound sub-diffraction limited gold nanoparticles (37.0 nm - 43.0 nm diameter) and cancer-associated fibroblast cells (CAFs) cultured over 7 days (**Methods**).

Our home-built RO-iSCAT system ^18^ involved a pair of galvanometer scanner to achieve customized off-axis oblique and azimuth (**Fig. 2a**). Two raw iSCAT image were recorded, one with (**Fig. 2b**) and one without gold nanoparticles (**Fig. 2c**, as background), and formed the final image iSCAT image after background is manually subtracted (**Fig. 2d**). Each raw image was recorded by positioning the galvanometer mirrors at a single azimuthal position. On the other hand, the RO- iSCAT image was captured after turning the galvanometer mirror at azimuthal angles 𝜑 from 0 to 360° continuously. To capture a full RO-iSCAT image, the camera exposure rate was synchronised to integrate over a series of oblique RO-iSCAT images over a single cycle of rotation, finally get Σ𝐼_scat_(𝜑) (acquisition speed up to 40 fps in our system). Here we demonstrated four RO-iSCAT images at each azimuth 0°, 90°, 180° and 270° (taken without rotational integration, **Fig. 2e**) and the full-integrated image (**Fig. 2f**). Considering that the final iSCAT and RO-iSCAT images were of the same field of view and taken over the same exposure time, the full RO-iSCAT images outputted significantly lower background noise and speckle than iSCAT background subtraction, as well as an alignment with our numerical simulation (inset of **Fig. 2f**).

**Fig. 2.**
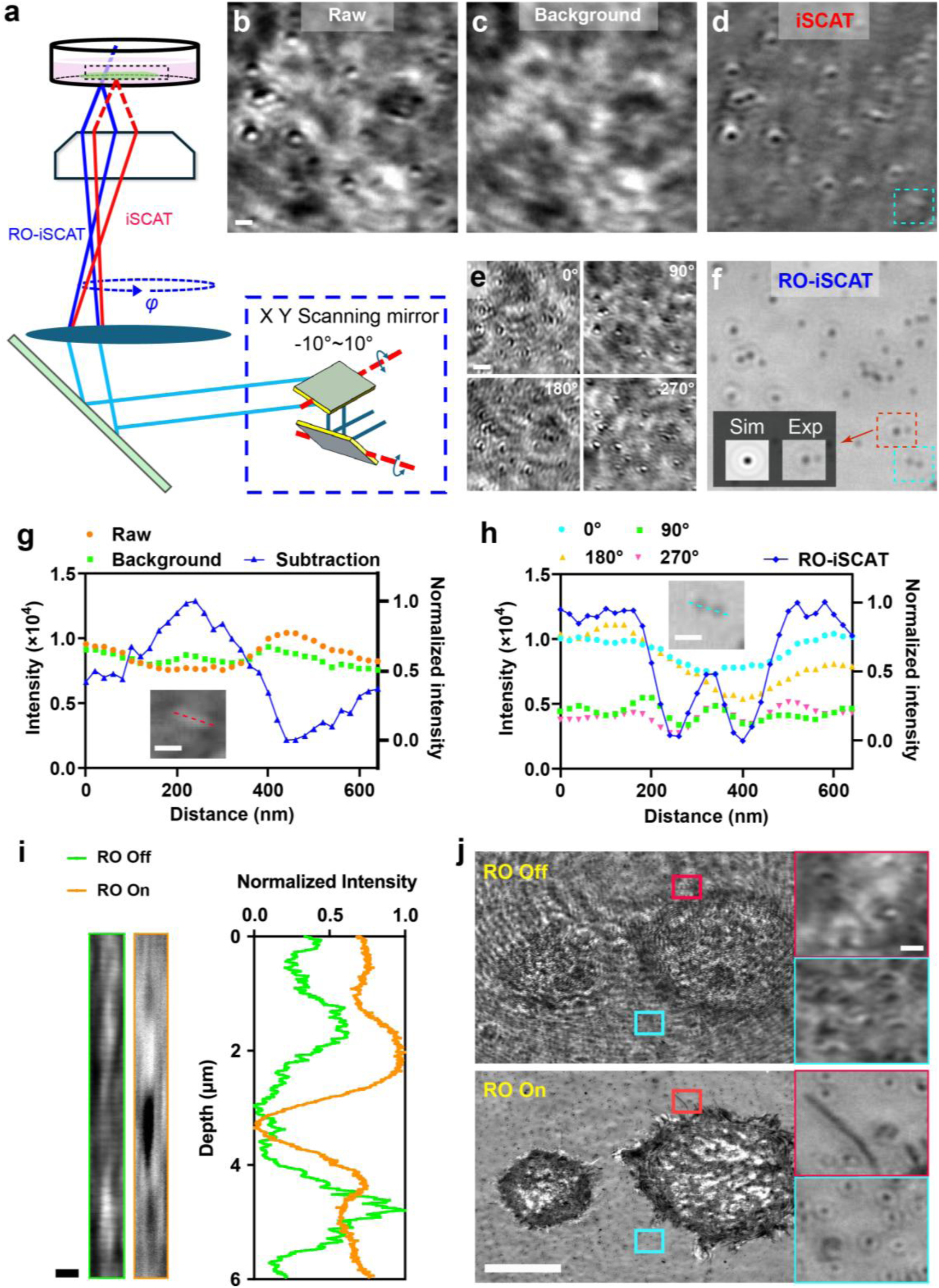
Reduction of speckle background through rotational integration. **a)** Diagrammatic sketch of conventional iSCAT and RO-iSCAT imaging method. Bottom right is the sketch map of orthogonal galvos and the transition from flipping at the galvo to lateral circling at the back focal plane, and finally to illumination with specific tilting angle and varying azimuth emitted from the objective lens. **b)** Raw iSCAT image of 40nm gold particles with 500 ms exposure time. Gold particles were dried thus adhered to the inner glass bottom, then PBS for 1:20 dilution to the initial storage liquid of particles. **c)** Manually selected background pattern from PBS-only control dish under the same exposure time. **d)** Final iSCAT image after post-processing of background subtraction. The dashed rectangle region highlights the blurring pattern in iSCAT but well-distinguished by RO-iSCAT. **e)** RO-iSCAT images from different incoming azimuths without integration. **f)** Final RO-iSCAT image with time-integrating during rotational scanning. **g)** Intensity profile of raw image, background and the final result after subtraction, along the dashed line in magnified image cropped from **d)**, the orange and background scatters are labeled by left y axis while the blue curve is by right y axis. **h)** Intensity profile of image at azimuth position of 0°, 90°, 180°, 270° and final integrated result along the **f)** region that corresponds to the dashed line, the blue curve is by right y axis while the scatters in other colours are labelled by left y axis. **i)** Z section and the corresponding profile along z axis from interference PSF captured from 40 nm particles, individually without rotational integration and with rotational integration configuration. **j)** Without rotational integration (top) against with rotational integration (bottom) modality of CAF cells seeded with 40 nm particles. **Scale bars**: **b)**-**d)**, **f)** 250 nm, **e)** 500 nm, **g)-i)** 400 nm, **j)** 5 μm.

To quantitate the improvement of RO-iSCAT over the background subtraction in iSCAT, we chose a smaller field of view (cyan dotted box in **Fig. 2d** and **2f**) and adopted the metric to determine if two closely spaced sub-diffraction limited gold nanoparticles can be resolved. From the line plot of intensity variation between adjacent nanoparticles (**Fig. 2g** and **2h**), it appeared that background subtraction and integration both possessed almost the same signal to noise ratio. However, rotational integration was able to fully resolve adjacent 40 nm particles, where separation between the 40 nm particles was approximately 170 nm that was close to Abbe diffraction limit of 163.7 nm, which was not possible using background subtraction for the same acquisition time (insets in **Fig. 2h** and **2g**). RO-iSCAT removes of the speckle noise in the background without any loss to intensity. Because no subtraction is made, the full dynamic range of the camera is preserved. Also we observed that there were spatially varying speckles intensities in the images taken from 𝐼_scat_(90°), 𝐼_scat_(270°) to 𝐼_scat_(0°) and 𝐼_scat_(180°) profiles which were uncorrelated and so were removed in RO-iSCAT Σ𝐼_scat_(𝜑) because of rotational integration (**Fig. 2e**).

To further validate our numerical model that rotational integration improves the imaging resolution along the axial direction (**Fig. 1e** and **Supplementary** Fig. 3), we captured the interferometric signal using a nano-stage that was moved along a fixed step interval of 10 nm. When comparing the axial intensity profiles (**Fig. 2i**), the axial intensity variation of RO-iSCAT shown higher contrast along the axial plane and matched well with Boundary-Element-Method simulation (**Supplementary** Fig. 3). Then, we evaluated the imaging performance of rotational integration with RO-iSCAT on membrane protrusion from adherent cells alongside with fixed 40 nm gold nanoparticles (**Fig. 2j**). While 40 nm gold nanoparticles (**Fig. 2j**, cyan boxed inset) were marginally visible, it was only RO-iSCAT that the fine membrane protrusions can be detected (**Fig. 2j**, red boxed inset). Moreover, the rotational integration effect can be more readily observed and quantified by discretizing the integration process (**Supplementary Video 5**) which illustrated that the increasing number of azimuthal scanning angles 𝜑 for integration will form a higher final SNR (**Supplementary** Fig. 5).

### Differing spatiotemporal dynamics between membrane trails, tethers and bridges

Benefit from the high SNR fringes, we put our focus on the protrusion growth and external connections. First, we captured a time-lapse dataset of endothelial cells with high dynamics (**Supplementary** Fig. 6, **Supplementary Video 6**). On the smooth cell membrane, multiple protrusion emerged in random directions, then converging toward another cell, with lamellipodia driving them aggregating toward the target location, ultimately connecting with the target cell. During the growth, the interference pattern on protrusion were varying, mainly the bright-dark periods (yellow arrows in **Supplementary** Fig. 6). Here we ask if RO-iSCAT interference patterns can be used to identify membrane types, particular cell-substrate versus membrane bridges between cells.

To answer this question, we examined RO-iSCAT images in a single culture of CAFs cells that is known to form tight networks i.e. fibrotic tissue. RO-iSCAT provided the clear FOV where a CAF cell adhered to glass coverslip and multiple CAF cells forming extensive membrane bridges over 20-30 µm long (**Fig. 3a** i and ii). We then selected three different membrane protrusion to characterise which were chosen based on their distinct types of interference patterns and their assumed spatial locations. Combining the lateral morphology (**Fig. 3b**) and relative axial position (**Supplementary** Fig. 7 and **Supplementary Video 7**), we formed a biological sketch (**Fig. 3c**) and shown the membrane protrusions far from the cell body exhibit uniform fringe intensity, indicating a flat height and membrane has adhered to the glass-bottom surface (Trail, Fig. 3b top). In contrast, protrusions between cell bodies (basal surface) display interference patterns with alternating bright and dark fringes, spaced approximately 0.5 micrometers apart. This suggested that the protrusion grows from the apical side slanting downward to the bottom surface with a rapid height gradient (Tether, **Fig. 3b** middle). Additionally, fringes with membrane bridges tend display interference patterns with alternating bright and dark fringes, spaced approximately 3 micrometers apart, forming connections at a similar height between two cells (Bridge, **Fig. 3b** bottom).

**Fig. 3.**
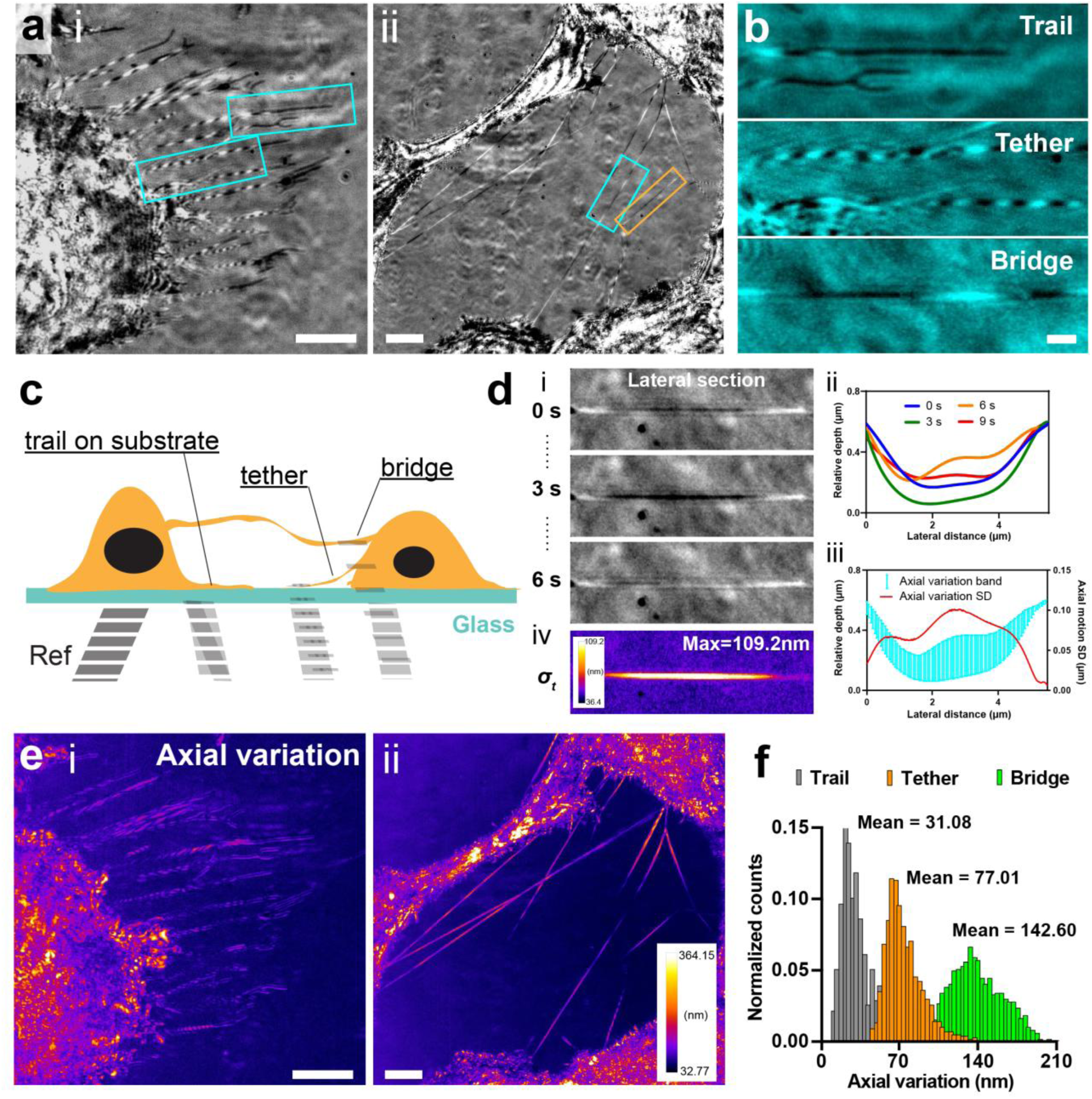
Identifying types of membrane trails and connections with RO-iSCAT. **a)** i and ii are raw RO-iSCAT of CAF cell membrane trails and connections from RO-iSCAT captured at 5 fps (50% duty cycle) over 10 seconds. **b)**, Magnified images of the cyan rectangle regions in **a)**. Top, cell membrane trails observed directly on substrates. Middle, cell membrane tethers mixed with membrane trails on substrates. Bottom, direct cell-cell tethers without any membrane trails on the surface. **c)** Proposed concept of biological diagrammatic sketch of neighbouring cells cultured on glass bottom dish, in which each main membrane protrusion type will create different scattering field. **d)** Procedure of calculating axial-variation map. i, RO-iSCAT fringe along one single protrusion at 0 s, 3 s and 6 s time points. ii, Axial distribution of this protrusion at several time points. Profiles were fitted from raw curve only for better presentation here (8^th^ degree polynomial with R^2^=0.94, 0.96, 0.92, 0.82 separately). iii, Standard deviation measuring the effective axial displacement at each pixel. iv. Pixel-level standard deviation on the entire 2D image relative depth over the time period. **e)** Calculated axial-variation maps of respective raw RO-iSCAT images in **a)**. **f)** Histograms counted from the trails and connections regions in **b)**, with mean values noted for membrane trail, tether and bridge groups. **Scale bars**: **a)**, **e)** 3 μm, **b)**, **d)** 500 nm.

Besides the morphology characteristic, we examined the motion modalities of these membrane protrusions from the entire time-lapse dataset by mapping from interference signal intensity to relative depth based on calibration data (**Sup.** Fig. 9, **Methods**). Kymograph is a classic tool for recording motion along one line over time (**Supplementary** Fig. 8), however, to capture the spatiotemporal changes across the whole imaging field, we applied a new axial-variation map (**Fig. 3d** i, ii, pixel-level standard deviation on the entire 2D image relative depth over the time period) to measure the effective range of axial displacement (**Fig. 3d** iii, iv). Because axial variation was applied across the whole imaging field, we can directly quantitate whole membrane protrusion dynamics directly from the intensity mapped to the magnitude of the axial variation (brighter intensity indicates larger axial fluctuations). To prove that this axial variation information was only retrieved using RO-iSCAT imaging, we also applied axial variation treatment to scattering-only images (**Supplementary** Fig. 8). It shown the distinct spatial temporal intensity changes were only observed under RO-iSCAT but not under scattering-only imaging (only up to 87 grayscale), indicating that this was a direct consequence of interferometry (**Supplementary Video 8**). The axial-variation map directly determined highly motile membrane bridges between CAF cells. The large range of intensity variation in RO-iSCAT images occurs in up to 300 nm z-axis movement along membrane bridge. This observation suggested that membrane bridges possess a taut behavior where axial movements are greater than lateral movements. Axial variation responses of the RO-iSCAT images (**Fig. 3e** i and ii) and the statistics of the values for all the three types of membrane protrusion (**Fig. 3f**) indicated a clear difference between the spatial temporal behavior among them. The mean values of the three grouped distributions histograms further illustrated that the suspended cell-cell bridges showed a 2-fold more axial movement (averaged range of 142.60 nm) than tethers (77.01 nm) and 4-fold more than trails (31.08 nm), even though they may physically appear tight and straight along.

### RO-iSCAT’s performance in tracking membrane protrusion

CAF cells can form extensive cellular networks that will be filled with various types of membrane protrusions particularly cell-cell membrane bridge ^27, 28^. Two separate cell co-culture experiments using CAF and KPC cells were adopted (**Methods**) here to track the formation and transition of protrusion and connections between identical and different cell types.

First, we examined whether the interferometric images from RO-iSCAT can track different membrane protrusions more effectively than that based on intensity-only morphological tracks in fluorescence imaging. We co-cultured two cell populations of CAF cells: wild type (WT) CAFs and CAF transfected with Lck10-GFP to label the plasma membrane (**Fig. 4a** i). Two separate cultures of WT and Lck10-GFP transfected CAF cells were grown over 3 days before both cultures were seeded into the same culture dish. The culture was imaged four hours after seeding under scattering-only (**Fig. 4a** ii), fluorescence (**Fig. 4a** iii), alongside with RO-iSCAT all under oblique highly inclined thin illumination (HiLO). A sequence of membrane protrusions (red dotted box in **Fig. 4a** iii) over ∼ 7.9 minutes, under HiLO fluorescence imaging (**Fig. 4b**) and RO-iSCAT (**Fig. 4c**), shown a transfected Lck10-GFP CAF cell protrusion its membrane towards neighboring WT- CAFs (not visible under fluorescence) imaging. Under RO-iSCAT imaging, we can readily observe both transfected-CAF and WT-CAF beginning to form membrane bridge ∼5µm length.

**Fig. 4.**
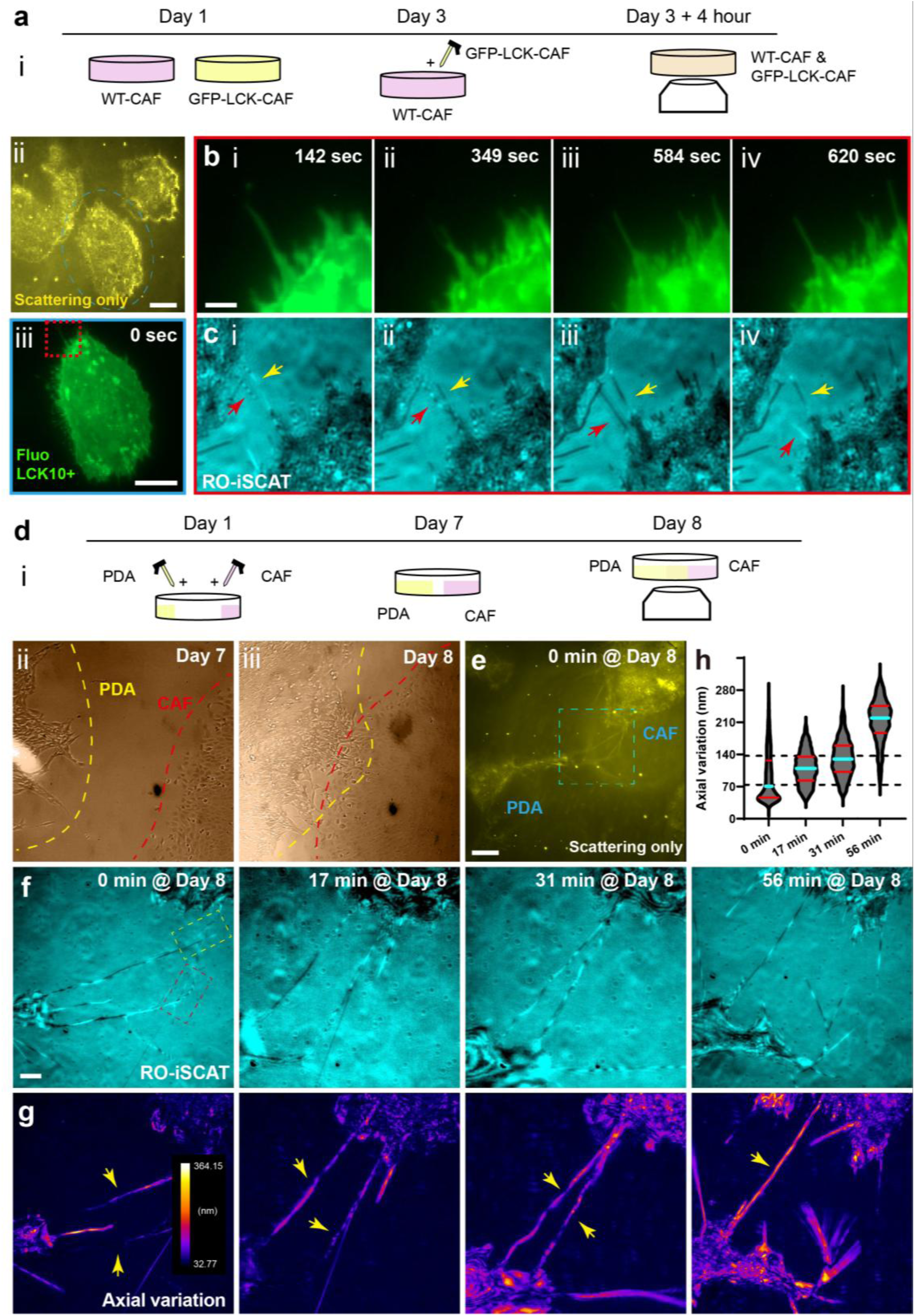
Tracking and quantifying protrusion between trails and connections with RO-iSCAT. **a)** i, Experiment time sequence of co-culturing Lck10-GFP transfected and non-transfected WT CAF cells. On day 1, WT CAF cells were seeded into 2 dishes and only one of them was transfected by Lck10-GFP dye. On day 3, we transferred the transfected cells into the WT CAF cell dish. 4 hours later, the mixed dish was uploaded for imaging. ii, Scattering-only image captured at 4^th^ hour on day 3 containing only one transfected cell (marked by light blue circle). iii, Magnified fluorescence image of the Lck10-GFP transfected cell in FOV (dashed light blue circle in ii. **b)** Time-lapse fluorescent images of the red rectangle region in **a)**. **c)** The comparative RO-iSCAT images where the two different arrows consistently track the same protrusion across different time points. **d)** i, Experiment time sequence of PDA and CAF cells co-culture. On day 1, we seeded CAF and PDA cells separately at opposite side of one dish. During day 1 to 8, each will migrate towards the center over 7 day-long culture in incubator. On day 8, imaging was performed where the two cell populations intersected. ii, iii, Bright field images under stereo microscope of PDA- CAF co-culture dish at day 7 and 8 after seeding. **e)** Scattering-only image of a FOV containing one PDA and CAF cell. **f)** Snapshots in time-lapse stack of the cyan rectangle region in **e)**, individually displaying the filopodia interacting, connecting, merging and the final merged dynamics. **g)** Axial variation map of **f)** counting from 50 frames (5 fps) around each time point. **h)** Violin plot of axial variation distribution along membrane protrusion in each time point. Cyan and red lines represent median and quartiles, respectively. Counted protrusions have been pointed out by yellow arrows. **Scale bars**, **a)**, **e)**, 5 μm, **b)**, **f)** 1 μm.

While fluorescence imaging permitted the identification of 2D morphology of the membrane protrusion, in contrast under RO-iSCAT, we observed a highly complex interference scattering patterns within the same protrusion. The intensity stripes changed from high to low spatial cycles (all arrows in **Fig. 4c** i, ii compared with **Fig. 4c** iii, iv). At time point 349 and 584 seconds, an adjacent stationary protrusion appeared to display a uniform dark stripe (yellow arrow in **Fig. 4c** ii and red arrow in **Fig. 4c** iii), suggesting the complete disassociation from the target cell individually. The variation of these intensity fringes indicated physical axial movements of the protrusion ^18^. Over 10 mins, these interferometric intensity patterns exhibit regular axial movement when associating and disassociating with neighboring cells (**Supplementary Video 9**). All these results suggest that RO-iSCAT overcomes the limit of fluorescence modality for the purpose of quantifying spatial temporal dynamics in 3D membrane protrusions. Besides these CAF-CAF intercellular connections, direct communication between CAFs and cancer cells mediated by surface receptors or adhesion molecules can play an much more important role in tumour progression ^29^. Hence, next we looked at whether CAF can form membrane bridges with pancreatic ductal adenocarcinoma (PDA) cells.

The second co-culture involved CAF and PDA cells from the murine KPC model, where we seeded the two cell groups separately at opposite side of the glass bottom dish where each will migrate towards the center over 7 day -long culture in incubator (**Fig. 4d** i). On day 7 and 8, we imaged the proximity of the cell (**Fig. 4d** ii and iii) under brightfield to observe the proximity of the 2 cells population indicated by yellow and red dotted line. On day 8, we used scattering-only images to identify the border between a CAF and PDA cell using cell morphology (**Fig. 4e**). Within the chosen field of view, we monitored the space between CAF and PDA cell (**Fig. 4e**, blue dotted box). RO-iSCAT imaging was conducted over the same field of view over 60 mins at 10 mins intervals (**Fig. 4f** and **Supplementary Video 10**). Using RO-iSCAT and interferometric signals, we identified active transition between cell-substrate connections to membrane bridges. Two individual connection appeared to gradually merge to one single protrusion through a twisting protrusion morphology ^10^. We can see that at the initial time point (0 min), the protrusion from a single CAF and PDA cell were first individually separated (yellow and red dashed box). Then at 17 min, the protrusion gradually merged to form direct membrane bridge. Starting from 31 min, the two separated membrane bridge merge through high degree of axial variation that maybe indicate large strain akin to twisting to form a single membrane connection at 56 min with a straighter morphology.

Whilst the interference intensity shown highly dynamics activities (**Supplementary Video 10**), it failed to reflect spatial temporal information visually. To measure the spatial temporal events, we applied axial-variation treatment to the RO-iSCAT images to measure fluctuation from the interference intensity signals (**Fig. 4g**), where higher value of axial variation indicates increasing protrusion motility. It directly indicated the increasing level of axial motion along each protrusion across each time point. Quantitatively, a violin plot (**Fig. 4h**) shows the time-variance distribution along the protrusion at each time point that indicates distinguishable mean values between membrane bridges forming between the PDA and CAF cells. Particularly, the spatiotemporal dynamics of the protrusion from tethers to formation of bridge between cells. Newly connected tether was observed at 17 min and 31 minutes displayed increasing axial motion up to 140 nm which was 2 times compared to initial protrusion tethers (0 min), and at the last 56 minutes, we observed a bridge was formed between the cells indicated by 210 nm range of axial motion (3-fold increase). This result indicates that RO-iSCAT interference signals may be used to identify spatiotemporal behavior of membrane bridges that is otherwise missed by classical 2D intensity- imaging (fluorescence or scattering-only) techniques.

## Discussion

In this study, we performed two investigations: 1) eliminate out-of-focus speckle background noise in RO-iSCAT interferometric signatures using rotational integration. 2) preliminary evidence that axial variation map of label-free RO-iSCAT images can measure and quantify spatial temporal membrane protrusion types that surpass conventional kymograph.

### Can RO-iSCAT operate with incoherent sources?

SNR of RO-iSCAT is defined by fringe visibility. Along the transverse and axial planes (x, y and z), fringe visibility is generally affected by the coherence of the light source and relative difference of both the optical path length and intensities between reflected and scattered intensities ^21^. Interferometric scattering (iSCAT) ^19, 20, 24, 30–32^ is gaining traction for label-free sub-cellular imaging due to its ability to detect nanoscale scatterers. RO-iSCAT could be treated as a partially coherent detection (temporal) along with the integration time of the camera. Out-of-focus interference fringes are suppressed based on the difference in path length. In supplementary, we have done some modeling of partially coherent sources indicating coherence plays a minimal role. We suspect that this rotational integral configuration can be of use for most interferometric microscopies.

### What are the key drawbacks of RO-iSCAT modality?

A common problem in interferometric imaging is the repeating interference fringes due to wrapped phase. This meant that direct 3D quantitative measurement of interference fringes in the axial plane becomes a challenge. Secondly, the axial iSCAT sections are taken with a moving objective lens or the first reflective surface of a glass coverslip which can incur additional phase shifts. This is a similar issue with common path interferometry. It will refresh background artifacts from optical components, and the moving length of the objective lens will not convert to the same change of interference intensity as described in the initial intensity-height relationship.

### To combine with optical tweezers to study membrane tension

Using interferometric spatiotemporal images of RO-iSCAT and in combination with optical tweezers ^33^, we expect to potentially quantify membrane tension of the twisting nascent filopodia ^10^ (**Fig. 4**). More recently, Belly et al ^34^ showed using optogenetics that 2D actin-driven protrusions can elicit rapid global membrane tension propagation resulting in long-range membrane flows. Using RO-iSCAT along with calibrated tweezers, one can study 3D axial membrane protrusion when dynamics forces (optical forces) are applied to the actin cortex.

### Incomplete abscission, phagocytosis and cell adhesion were quantified by measuring cytoplasmic bridge properties, 3D filopodial dynamics on bacteria and nanotopologies

The physical properties of membrane bridges provide insight into abscission completion during the final stages of cell division ^35^, and 3D filopodia extensions define the distinct stages of phagocytosis as immune cells clearing bacteria ^4^and recognition of nanotopologies that guides cell migration ^7^. While many studies utilize confocal microscopy, its axial resolution and high phototoxicity remains a key limiting factor for live axial imaging and tracking of filopodia. ROiSCAT’s value lies in its axial sensitivity and low phototoxicity, which operates below the diffraction limit and require minimal power (∼microwatts). Consequently, we anticipate that ROiSCAT will be highly valuable in quantifying incomplete abscission, cell migrating on nano- scale surfaces and phagocytosis.

## Conclusion

We showed that RO-iSCAT interference patterns generate highly distinctive spatial-temporal interference intensity patterns between different cell membrane protrusion, i.e. membrane that are tethered onto substrates, trails (e.g. migrasomes) and membrane bridges between adherent cells across large physical gaps ^27, 36^ Unlike scattered only or fluorescence signals used in kymograph, the spatial temporal interference patterns created unique axial variation plots for image-based classification. These characteristics were shown to be applicable across a range of adherent cell types, including endothelial, CAF and PDA cells. This pilot study has highlighted the potential of our method in extracting membrane specific interferometric patterns that eludes fluorescent imaging. This study enables the classification of membrane protrusions that, despite possessing identical chemical compositions, are differentiated by their interactions, thus offering a qualitative comparison of cell-cell communication at the nanoscale in living cells

## Author Contributions

Conceived project and directed research: W.M.L.

Prepared samples: Y.J.L., P.T., W.M.L., D.H., T.G.P, P.T.

Built model and programed simulations: J.L., M.G., H.L.

Designed and performed experiments: J.L., Y.J.L., W.M.L.

Examined and analyzed data: J.L., W.M.L.

Wrote manuscript: J.L., W.M.L., with advice from all authors.

Provided biological insight and advice: Y.J.L., W.M.L.

Supervised research: M.G., H.L., W.M.L.

## Acknowledgments

We thank Alpha Yap (IMB, UQ) and Melanie White (IMB, UQ) on membrane protrusion and filopodia discussions; and Hari Shroff (Janelia, HHMI) on the RO-iSCAT. We acknowledge the Australian Research Council (DE160100843, DP190100039, and DP200100364) and NHMRC (APP2000485) for their support. H.L. acknowledges funding from the National Natural Science Foundation of China (62427807) and the Talent Program of Zhejiang Province (2021R51004). M.G. acknowledges funding from the National Natural Science Foundation of China (62475232). J.L. acknowledges the support from Zhejiang University Global Partnership Fund.

## Methods

### Experiment setup

#### RO-iSCAT/Scattering data acquisition

Our rotating optical coherent scattering platform (ROCS) is equipped with 60× 1.49NA oil immersion objective lens (Olympus) and an sCMOS camera (PCO edge 4.2) for wide field illumination and detection resulting in a pixel size of 20 nm and a full field-of-view over 41 microns × 41 microns.

A 488 nm laser beam is directed onto a two-axis galvanometer and conjugated onto the back focal plane of the objective lens to generate an oblique angle and rotational azimuth. The camera performs capture under a preset framerate (up to 100 fps) and duty cycle while the incident beam separately rotates at a fixed speed of 200 rounds per second. ROCS sets oblique illumination angles to switch between interferometric (22°) and scattering imaging (60°) modes for simultaneous multimodal imaging. In addition, RO-iSCAT modality requires a glass bottom culture dish or coverglass for generating reference reflection light, so in the other scattering mode, the reflection will be rejected by an electronic amplitude filter (diaphragm) placed at the imaging back focal plane.

#### iSCAT data acquisition and post-processing

The iSCAT raw images were acquired from ROCS system by fixing the galvanometer with 𝜃 = 22° and 𝜙 = 0° under RO-iSCAT mode. Because ROCS platform adopts the strategy that moving the objective lens or sample container to adjust the focal plane, the reference field is different at different focal positions thus caused varying background images. A series of background images were acquired from an empty glass dish or coverglass under different focal positions and exposure times.

For isolating pure background artifacts arising from optical components, we meticulously maintained the system’s condition, particularly the stage position, and replaced the petri dish with another one only with PBS as a control, then captured the background image (**Fig. 2c**). Noted that the apparent fringes in background will blur or focus and undergo overall intensity changes during the stage moving along the z-axis. Thus, background subtraction necessitates the 1) pre-collection of a series of background patterns at each z position and after each biological acquisition, 2) a manual selection for matched background due to the limited repeatability of the translation stage. After background subtraction (**Fig. 2d**), the interference image marginally excluded some ambiguous artifacts, but generally, no new fringes of gold particles emerged from the background.

#### Calibration protocol

We used 40nm AU nanoparticles in a cell culture dish full of DMEM solution to illustrate the sinusoidal relationship between interferometric intensity and depth gradient (**Supplementary** Fig. 9a, **9b**) in RO-iSCAT. The exposure time of the camera was set to 85ms.

The dish was placed on a piezo nanostage that adopted axial sweeping of the sample across the focus at 10 nm steps. The axial intensity map of a single nanoparticle is plotted as the orange scattered plot, and the black line indicates the moving average of 8 points (**Supplementary** Fig. 9c). The cursors represent the linear region that can be used to map from intensity (ΔI) to axial displacement (Δz).

### RO-iSCAT model

#### Field model from physical optics theory

In RO-iSCAT, particles of a sample are illuminated by incident coherent laser filed 𝐄_inc_propagating from the objective lens and create a scattering field 𝐄_scat_. Meanwhile, the glass bottom of petri dish or coverglass reflects part of the incident light and form a weaker reflection field 𝐄_refl_. The reflection field 𝐄_refl_ and scattering field 𝐄_scat_ return to the imaging plane and jointly form the interference pattern. ^37, 38^

The final fields reaching the camera are the convolution of the initial field in object space and the intrinsic coherent transfer function 𝐂 of the optical system, and we get the interference pattern as

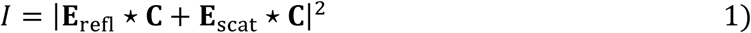

We built the Cartesian coordinate system *xyz* where z-axis is fully aligned with the optical axis of the objective lens, the coverglass and the focal plane of the objective lens are respectively set as 𝑧 = 0 and 𝑧 = 𝑧_𝑓_. Considering that nano-scale system usually adopts high-NA design, we assume that 𝐂 is a vector with an impulse response function as amplitude for simplicity, correspondingly, the interference pattern can be treated as formed by the initial reflection and scattering field reaching the focal plane 𝑧 = 𝑧_𝑓_

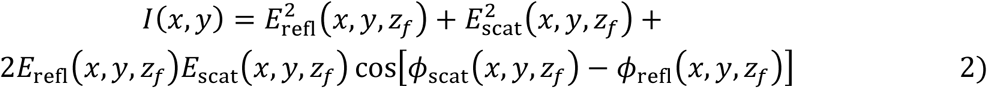

We assumed the laser beam with constant intensity in section instead of Gaussian distribution, and an incident at the oblique angle 𝜃 to the z axis and rotational azimuth 𝜑 around it. Ignoring the 𝜔𝑡 item that describes how the wave evolves over time and supposing the initial phase at (0,0,0) as zero

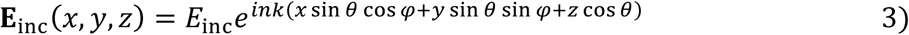

where 𝑛 is the refractive index of the air and 𝑘 = 2𝜋⁄𝜆 is the vacuum wavevector. As for the reflection field from oblique incidence, the amplitude and phase changes are complexly determined by the oblique angle according to the Fresnel formula, so we simply noted reflective index 𝜏_𝜃_ and phase shift 𝜙_𝜃_ as function of 𝜃 in reflection field

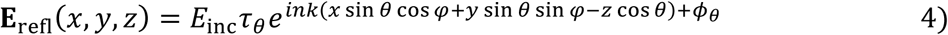

We consider a single nano-scale particle with sub-wavelength size and 𝑛_𝑝_ density located at (𝑥_𝑝_, 𝑦_𝑝_, 𝑧_𝑝_). The laser is scattered by the particle with the initial phase

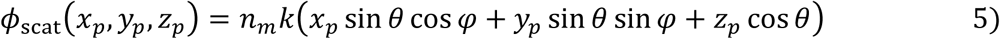

and amplitude variation to

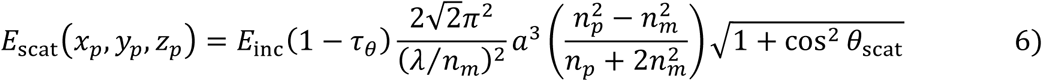

in which 𝑎 is the particle radius and 𝜃_scat_ is the scattering angle for representing anisotropic scattering efficiency. By combining the initial phase and amplitude, we can achieve the scattering field with spherical wavefront

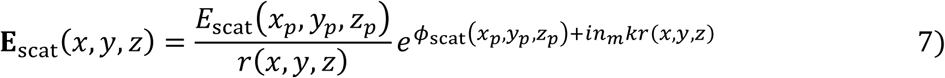

where 𝑟 is the propagation length 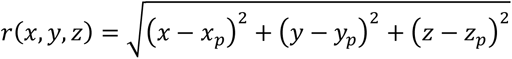 and cos 𝜃_scat_ = (𝑧_𝑝_ − 𝑧_𝑓_)⁄𝑟(𝑥, 𝑦, 𝑧).

And finally, we obtain the constant, amplitude, and phase items of the interference field at the focal plane

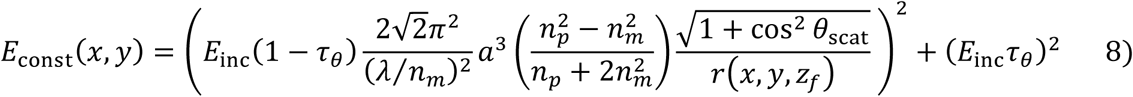

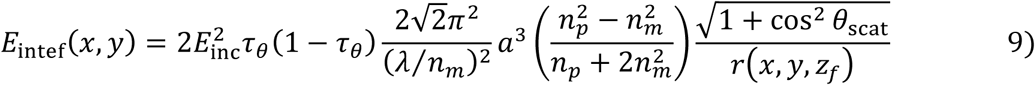

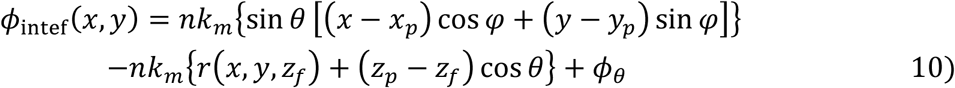

The equation of the interference phase relates to two key series of variables, (𝜃, 𝜑) for describing the incidence off principle optical axis and 𝑧_𝑝_ − 𝑧_𝑓_ for measuring the defocused length. 𝑧_𝑓_ is typically maintained as we usually fix the relative position between objective lens and container during imaging process. iPSF can be generated by changing 𝑧_𝑝_ − 𝑧_𝑓_ and 𝑧_𝑓_, respectively corresponding to the situation axial movement of 1) sample-only and 2) focal plane by adjusting the objective lens or the whole sample container (**Supplementary** Fig. 6). To visualize their roles in phase difference, we correspondingly split it into one off-axis and one defocused item

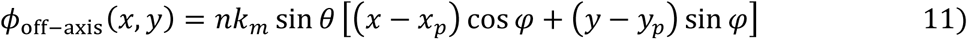

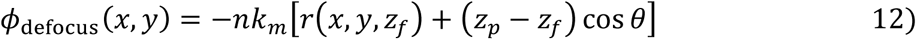

#### Lateral shift of out-of-focus pattern

In simulation, we set 𝜃 to 22° as a constant while 𝜑 in the range of 0°∼360° as a variable considering the configuration of our RO-iSCAT and place the phantom particle at the center of FOV (0,0, 𝑧_𝑝_). The container position was fixed for a constant 𝑧_𝑓_and we investigated the field created by defocused particle at 𝑧_𝑝_ with Δ𝑧 = 𝑧_𝑝_ − 𝑧_𝑓_ defocused length.

To mathematically quantify the lateral shift of pattern, we took the partial derivative of the total phase difference respectively to 𝑥, 𝑦

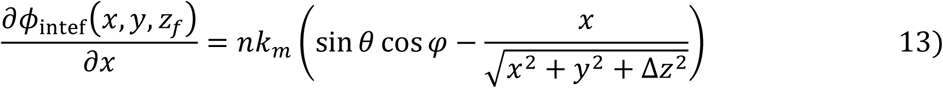

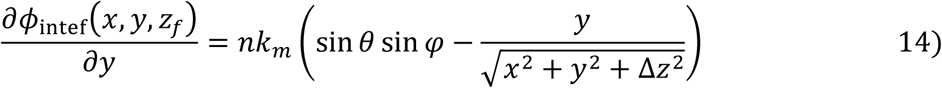

From which we can achieve the specific extremum point (𝑥_𝑒_, 𝑦_𝑒_, 𝑧_𝑝_)by setting the gradient simultaneously to 0

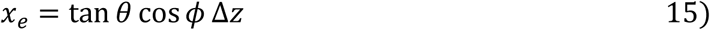

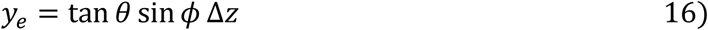

and the distance of its biased to centre point is

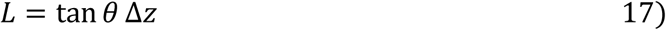

### Quantitative image quality analysis

#### Measuring the radius of lateral shifting

The focused position of 2 microns markers was set as zero axial position and gradually moved the along z-axis from negative 2000 nm to positive 2000 nm in 10 nm step using a high-dynamics Z nano-positioning stage (Physikinstrumente P-736.ZR1).

At each axial position, the camera got images from 12 azimuth points. The 12 centers of the marker were labeled and recorded by Manual-tracking plugin in Fiji (ImageJ2 core), then fitted the circle from the shifted centers to get the radius. We used negative and positive signs to distinguish from clockwise and counterclockwise rotational shifting.

#### Interference fringe contrast for azimuth-sampling dataset

The RO-iSCAT images were used as ground truth, i.e. the pure signal, while the middle outputs as the polluted overlay with noise and artifacts. During azimuth down sampling, due to lateral shifting direction varying at different azimuth point, the asymmetric azimuth combination (i.e., odd number sampling points) will retain some of this property. This results in a slightly distorted morphology that does not perfectly match the ground truth. Therefore, instead of subtracting the pure signal to determine the noise level, a common approach in SNR calculations, we directly use the down azimuthal sampling image to measure the level of noise.

We respectively selected 8 rois from the empty background and signal-intensive cell regions, then valued the pollution of messy noise to smooth structures by

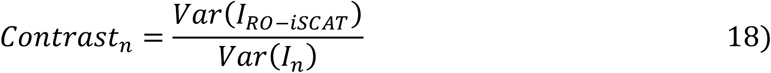

#### Simulation and SNR-measurement for speckle-noise

According to experimental observation (**Supplementary Video 1**), speckle noise is frame- uncorrelated and reserves the property of lateral shift same as physical particles. Hence, we model them as random particles with varying reflectivity value at one out-of-focus plane that generate unexpected fringes onto focal plane. For **Fig. 1e**, in a 200×200 pixels (4μm×4μm) FOV, we uniformly set one single particle on focal plane alongside with 600 particles at 1 μm depth (speckle noise from out-of-focus plane), then overlap all the fringes simulated from our model. Specifically, while the reflectivity of in-focus particle is set to 1 as reference, the relative reflectivity among those 600 speckles follows the Gaussian distribution but with an absolute operation to avoid negative values |𝒩(0, 𝜎^2^)|

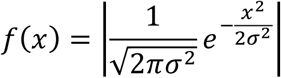

where we use 𝜎 to measure the speckle noise level.

To calculate the SNR of the synthetic fringes under iSCAT and RO-iSCAT modality, we selected the central region 𝐼_𝑠_ (5×5 pixels) from the on-focus particle fringe to calculate the variance of pure signal. As to the variance of speckle noise, we measure the variance of the overlapped fringe (on-focus particles and out-of-focus speckle) excluding the central region, 𝐼_𝑛𝑜𝑖𝑠𝑒_.

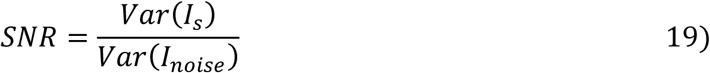

### Sample preparation

#### Cell culture

All reagents for cell culture were sourced from Thermo Fisher Scientific (Waltham, MA, USA). CAF and PDA cells (Passage 30) were maintained in T75 flasks with DMEM supplemented with high glucose (4.5 g/L), 10% fetal bovine serum, L-glutamine (4 mM) and pyruvate (1 mM) at 37°C and 5% CO2. Cells were split 1:20 at 80% confluence. Primary human lung microvascular cells (HMVEC, Lonza) were cultured in EGM2-MV2 Bulletkit (Lonza) at 37°C and 5% CO2 and split at a 1:6 ratio at 80% confluence.

#### Live cell imaging

Cells were first grown to 80% confluence and then 100,000 cells seeded to 29 mm glass bottom dishes (#1 coverslip, CellVis). Cells were incubated at 37°C and 5% CO2 for at least 1 hour prior to imaging.

#### CAF cell transfection

For fluorescent labeling of the CAF cell membrane, we used an Lck10-GFP plasmid (generous gift from the late Katharina Gaus). The Lck10-GFP consists of the first 10 amino acids of the membrane protein Lck and eGFP linked to the C-terminus. Plasmids were propagated in E.coli in LB Medium and purified using a miniprep kit (Genejet, Thermo Scientific). CAF cells were transfected with polyethyleneimine (40 kDA, linear, Polysciences Inc) using 9 ug PEI and 3 µg DNA per 29 mm dish over 48 hours prior to imaging.

#### Nanoparticles

Gold nanoparticles (40 nm) were diluted 1000 times in distilled water. 1 mL of diluted nanoparticles were then added to a dry 29 mm glass bottom dishes (#1 coverslip thickness, CellVis) and allowed to dry for 1 hour at room temperature. Prior to imaging, 1 mL of 1X PBS was carefully pipetted along the walls of the dish. The dish was then mounted onto a Z nano-positioning stage (Physikinstrumente P-736.ZR1) for imaging.

#### CAF cell with nanoparticle

40 nm gold nanoparticles were dried onto a glass bottom dishes (#1 coverslip thickness, CellVis) for 1 hour as described above. The dried particles were then immersed in high glucose DMEM prewarmed to 37°C. 1 mL of CAF cells (100,000 cells/mL) was then added dropwise onto the dish and incubated for 1 hour at 37°C and 5% CO2 and then mounted onto a heated stage for imaging.

#### CAF cell co-culture (WT and Lck10)

We seeded 100,000 cells into two separate 29 mm glass bottom dishes (#1 coverslip, CellVis). After 24 hours, one dish was transfected with Lck10-GFP plasmid and incubated for another 48 hours. Cells in WT and transfected dishes were then detatched with trypsin (0.25% (w/v)) and EDTA (1 mM) and 50,000 cells each seeded and mixed into a new 29 mm glass bottom dish. Cells were incubated for 4 hours at 37°C and 5% CO2 prior to imaging.

#### CAF and PDA co-culture

To create separate CAF and PDA cell populations on a single dish, cells were concentrated to 1 million cells/mL and 100 μL were pipetted to separate corners of a 29 mm glass bottom dish. Cells were incubated for 1 hour 37°C and 5% CO2 to allow the cells to attach to the glass dish. Cell attachment was monitored under a brightfield microscope. Cells were then supplementing with 1 mL of DMEM and incubated for 7 days, with media changed every 2 days.

## Data availability

The data that support the findings of this study are included in **Figs. 1–4**, **Supplementary** Figs. 1–10 and **Supplementary Videos 1–10**. All experimental data from figures (**Figs. 2**, **3**, **Supplementary** Figs. 5-9) are publicly available at https://doi.org/10.5281/zenodo.14960905. Other time-lapse co-culture datasets (**Fig. 4**) are available from the corresponding author W.M.L upon request due to their large file size.

## Code availability

All numerical modelling and analysis were achieved using Python 3.11.0. Generation of iPSF by Boundary-Element-Method was performed in MATLAB (Mathworks, R2022b). Customized RO- iSCAT model and analysis codes are available at https://github.com/ejunyuliu/RO-iSCAT. Initial numerical iSCAT model was installed from https://github.com/manoharan-lab/applied-optics-iscat-code. iSCAT software based on Boundary-Element-Method platform was downloaded from https://pubs.acs.org/doi/suppl/10.1021/acsphotonics.4c00621/suppl_file/ph4c00621_si_001.zip.

