## Supplementary info for "Using rotational integration of oblique interferometric scattering (RO-iSCAT) to track axial spatiotemporal responses of membrane protrusions"

**Supplementary Note 1**
Numerical simulation of RO-iSCAT

**Supplementary Figure 1**
Numerical simulation of RO-iSCAT lateral 2D iPSF
**Supplementary Figure 2**
Transforming RO configuration to conventional PSF
**Supplementary Figure 3**
3D iPSF comparison between iSCAT and RO-iSCAT
**Supplementary Figure 4**
Interference fringe SNR as a function of out-of-focus speckle noise level, comparing iSCAT and RO-iSCAT
**Supplementary Figure 5**
Reduction of speckle background through rotational integration
**Supplementary Figure 6**
Filopodium growth and external connections in endothelial cells
**Supplementary Figure 7**
Relative axial position of membrane tethers and bridges
**Supplementary Figure 8**
Comparing intensity variation on membrane protrusion under RO-iSCAT versus scattering-only modality
**Supplementary Figure 9**
Calibrating the intensity and axial displacement with 40nm nanoparticles
**Supplementary Figure 10**
iPSF comparison between two axial moving ways

**Supplementary Video 1**

Shifting speckle pattern under azimuthal scanning. Sample are 40nm gold nanoparticles on coverslips.

**Supplementary Video 2**

Simulation of interference pattern at 2 different axial planes; 10 nm and 10 μm of a single particle. Oblique illumination of 22° along multiple azimuthal angles (0 ° -360 °) along with defocused phase and resulting interference pattern.

**Supplementary Video 3**

Rotational scanning without and with integration with dried 2 μm size particles on glass slide.

**Supplementary Video 4**

Modelling different interference point spread function with widefield, confocal, rotational and rotational oblique.

**Supplementary Video 5**

Comparison without and with rotational integrated images of a single adherent cancer associated fibroblasts (CAFs).

**Supplementary Video 6**

Time-lapse of CAF cells (LCK-10 GFP+) in co-culture under Highly Inclined Laminated Plane (HiLO) fluorescence and RO-iSCAT.

**Supplementary Video 7**

Sequence of axially stepped RO-iSCAT imaging of CAF and KPC cells forming multiple membrane protrusions.

**Supplementary Video 8**

Comparison of scattering-only and RO-iSCAT of CAF cells co-culture along different axial planes

**Supplementary Video 9**

Time-lapse of scattering only and RO-iSCAT imaging of CAF cells with different types of membrane protrusions (trails, tethers, bridges)

**Supplementary Note 1: Numerical simulation of RO-iSCAT**

We built the xyz coordinate system where z-axis is fully aligned with the optical axis of objective lens, the coverglass and focal plane of objective lens are respectively set as $z = 0$ and $z = z_f$. Considering that nano-scale system usually adopts high-NA design, we assume that **C** is a vector with an impulse response function as amplitude for simplicity, correspondingly, interference pattern can be treated as formed by the initial reflection and scattering field reaching at the focal plane $z = z_f$. We derivated the field model for rotational-oblique incident interferometric scattering (see **Methods**) and got the final interference phase:

$$\phi_{\text{intef}}(x, y) = nk_m\{\sin\theta\,[(x - x_p)\cos\varphi + (y - y_p)\sin\varphi]\} \\ -nk_m\{r(x, y, z_f) + (z_p - z_f)\cos\theta\} + \phi_\theta \qquad 1)$$

where the $n$ is the refractive index of the air and $k = 2\pi/\lambda$ is the vacuum wavevector, in addition, the beam incident at the oblique angle $\theta$ to the z axis and rotational azimuth $\varphi$ around it.

The equation of the interference phase relates to two key series of variables, $(\theta, \varphi)$ for describing the incidence off principle optical axis and $z_p - z_f$ for measuring the defocused length. $z_f$ is typically maintained as we usually fix the relative position between objective lens and container during imaging process. To visualize their roles in phase difference, we correspondingly split it into one off-axis and one defocused item

$$\phi_{\text{off-axis}}(x, y) = nk_m \sin\theta\,[(x - x_p)\cos\varphi + (y - y_p)\sin\varphi] \qquad 2)$$

$$\phi_{\text{defocus}}(x, y) = -nk_m[r(x, y, z_f) + (z_p - z_f)\cos\theta] \qquad 3)$$

In simulation, we set $\theta$ to 22° as a constant while $\varphi$ in the range of 0°~360° as a variable considering the configuration of our RO-iSCAT and place the phantom particle at the center of FOV $(0,0,z_p)$. The container position was fixed for a constant $z_f$ and we investigated the field created by defocused particle at $z_p$ with $\Delta z = z_p - z_f$ defocused length.

Azimuth position $\varphi$ determined the direction of gradient in off-axis item (**Supplementary Fig. 1** first column) while $\Delta z$ broadens the pattern (reduced the gradient) in defocused item like PSF (**Supplementary Fig. 1** second column). Since they are not all centrally symmetrical, the total phase difference will perform an extremum at a specific position, which will be much more far away from the centre under a large defocusing length 10 microns (**Supplementary Fig. 1b**) than 10 nanometres (**Supplementary Fig. 1a**). To mathematically quantify this kind of lateral shift of pattern, we deviated the distance of its biased to centre point (see **Methods**) is

$$L = \tan\theta\,\Delta z \qquad 4)$$

which is surprisingly simple but suggests that rotating oblique illumination will create corresponding interference pattern in the plane perpendicular to the laser. This tilting defocused pattern will be projected onto the focal plane along the incident direction, thus results in lateral shifting and frequency changes to the interference rings both along the direction orthogonal to the azimuth. During rotational illumination, i.e. 0° ~ 360° azimuth, the pattern will exhibit circumferential lateral-shifting with the radius determined by defocused length (**Supplementary Fig. 1**, **Supplementary Video 2**).

We verified this relationship on experimental imaging on approximately 2 microns beads initially at focusing plane but gradually moved along z-axis from negative 2000 nm to positive 2000 nm in 10 nm step using a high-dynamics Z nano-positioning stage (Physikinstrumente P-736.ZR1) (**Supplementary Video 3** left). When translating from negative defocusing length to positive, the lateral shifting weakened first until reaching the focused position, after which it was going to strengthen. Additionally, we manually measured the length of shifting for the scatter plot (**Fig. 1c**) that illustrates a nearly matched slope that confirms the linear correlation between the radius of lateral shifting and the defocused length. Benefit from the varying lateral shifting, valid focusing information survives due to the scarcely shifting while the defocused background and noise will be averaged out by integrating the patterns with shifts along different azimuths (**Supplementary Video 3** right, **Supplementary Video 5**).

### Supplementary Figures

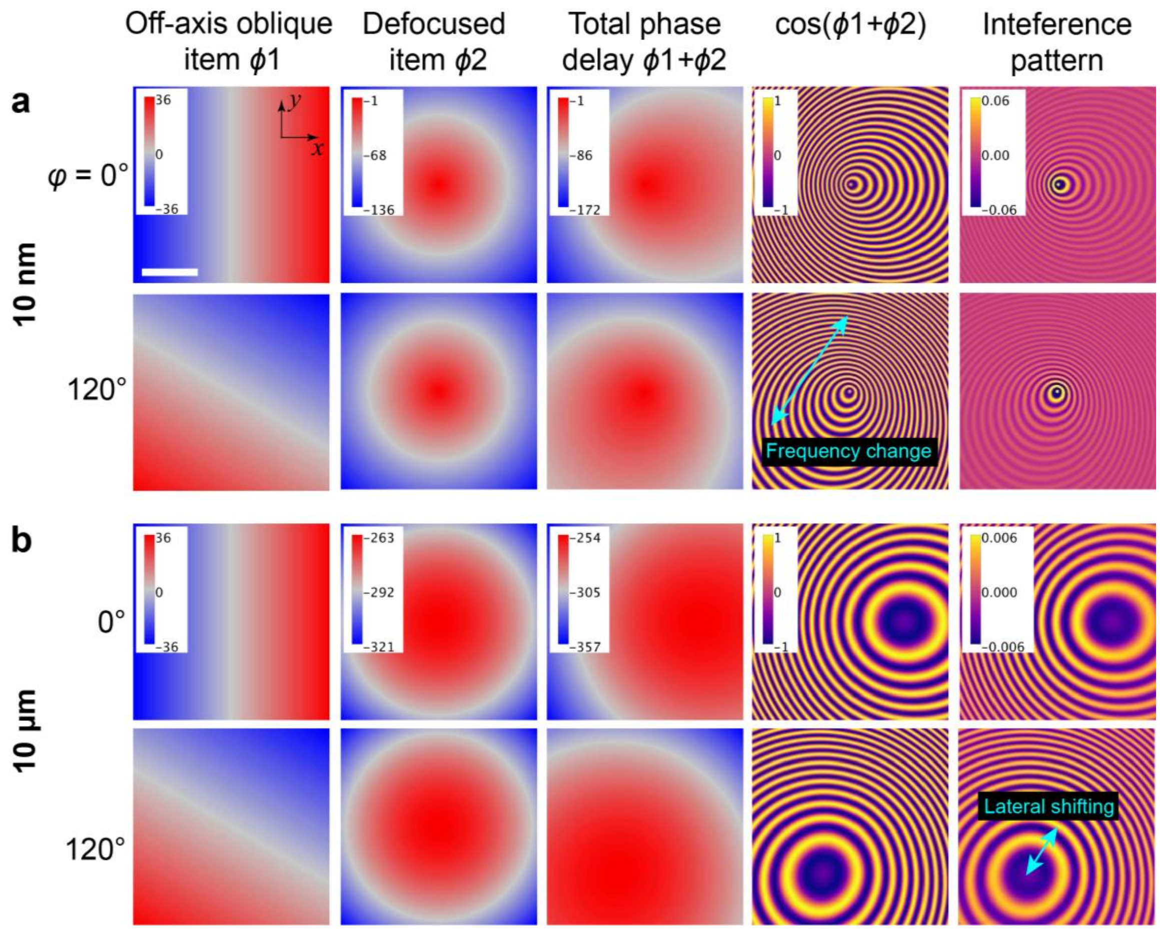


**Supplementary Fig. 1 Numerical simulation of RO-iSCAT lateral iPSF. a)** Numerical simulation with 14 microns FOV and 50 nm step based on the modelling. Including off-axis, defocused, total phase difference and interference pattern at 0°/120° with a 10 nm defocused length respectively. The cyan arrow highlights the frequency change of interference rings along the direction orthogonal to azimuth. **b)**, Phase difference and interference pattern at 10 microns defocused length. The cyan arrow indicates a larger radius of lateral shifting. **Scale bars**: **a)** 3 μm.

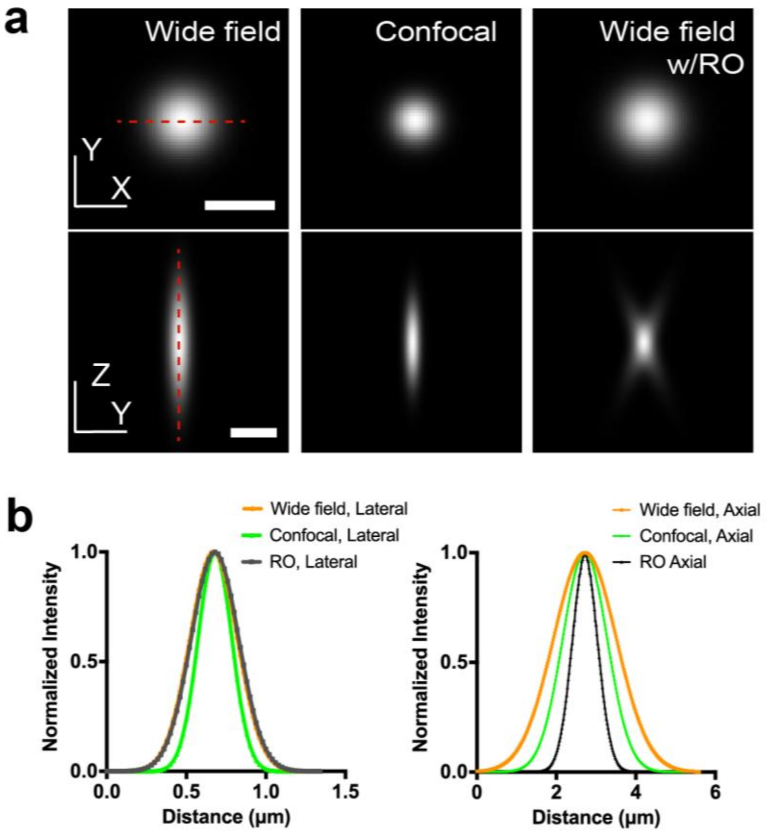


**Supplementary Fig. 2 Transforming RO configuration to conventional PSF. a)** Simulated XY and ZY section of wide-field PSF (left), confocal PSF (middle) and wide-field PSF with rotational-oblique configuration (right). All under 0.8 NA objective lens. **b)** Profile of the 3 kinds of PSF along the red dashed line in **a)**. **Scale bars**: **a)** 0.5 μm (top) and 1 μm (bottom).

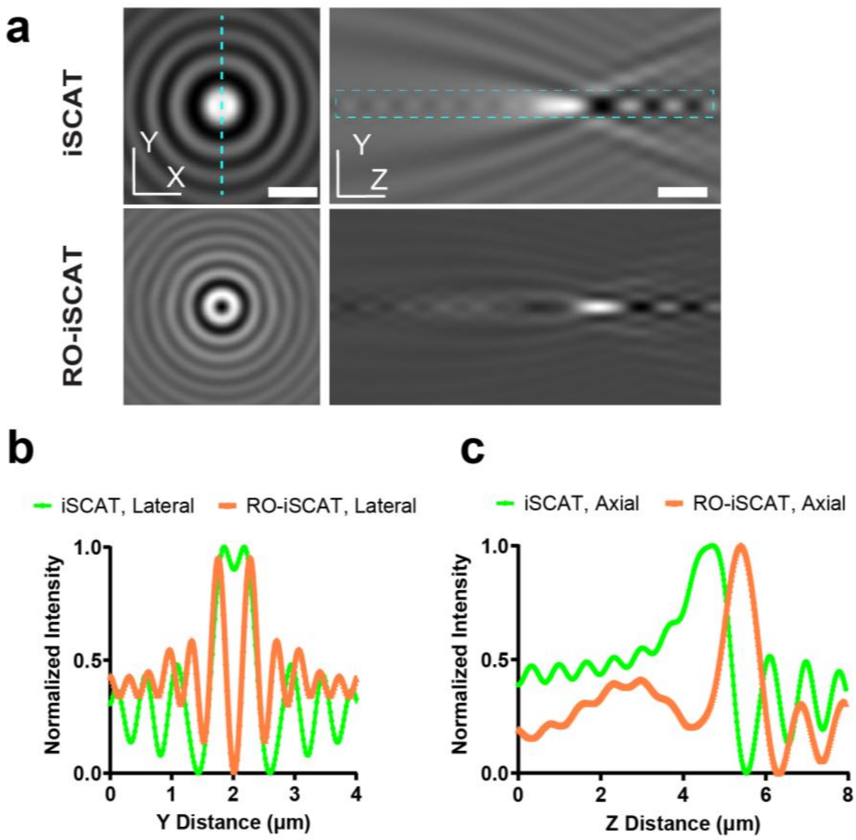


**Supplementary Fig. 3 Three-dimensional iPSF comparison between iSCAT and RO-iSCAT. a)** Simulated XY and ZY section of interference PSF, individually from wide-field, and rotational-oblique configuration. **b)** Profile of lateral interference PSF in focal plane, i.e. along the cyan dashed line in XY section. **c)** Profile of the axial interference PSF along the width of the cyan dashed rectangle in the YZ section, averaged over the height of the rectangle. **Scale bars**: **a)** 1 μm.

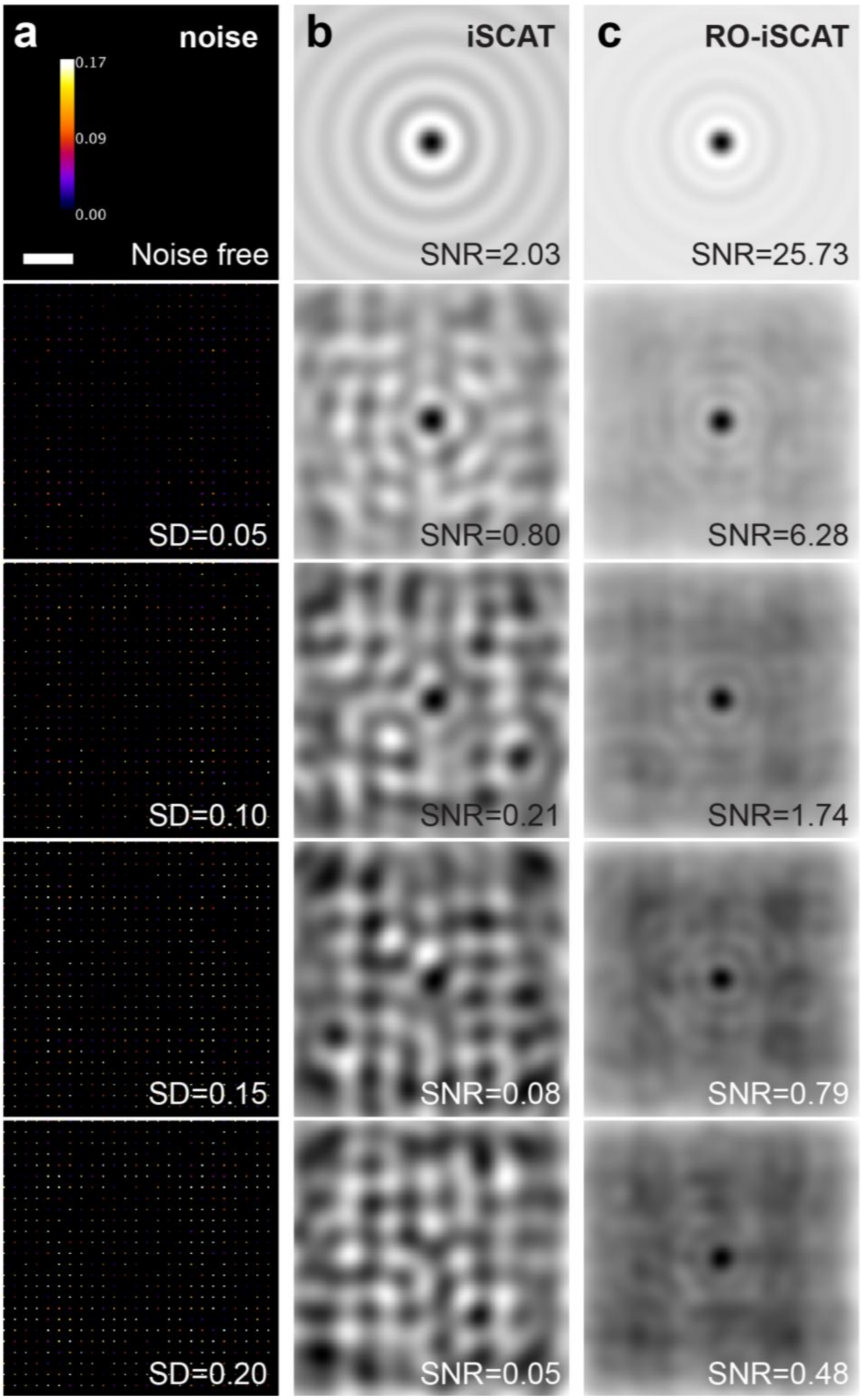


**Supplementary Fig. 4 Interference fringe SNR as a function of out-of-focus speckle noise level, comparing iSCAT and RO-iSCAT**. **a)** Input speckle noise map at 1-micron depth. Colormap refers to the relative reflectivity value of speckles compared with in-focus signal particle. **b)**, **c)** iSCAT versus RO-iSCAT fringes under corresponding speckle noise level. **Scale bars**: **a)-c)** 0.5 μm.

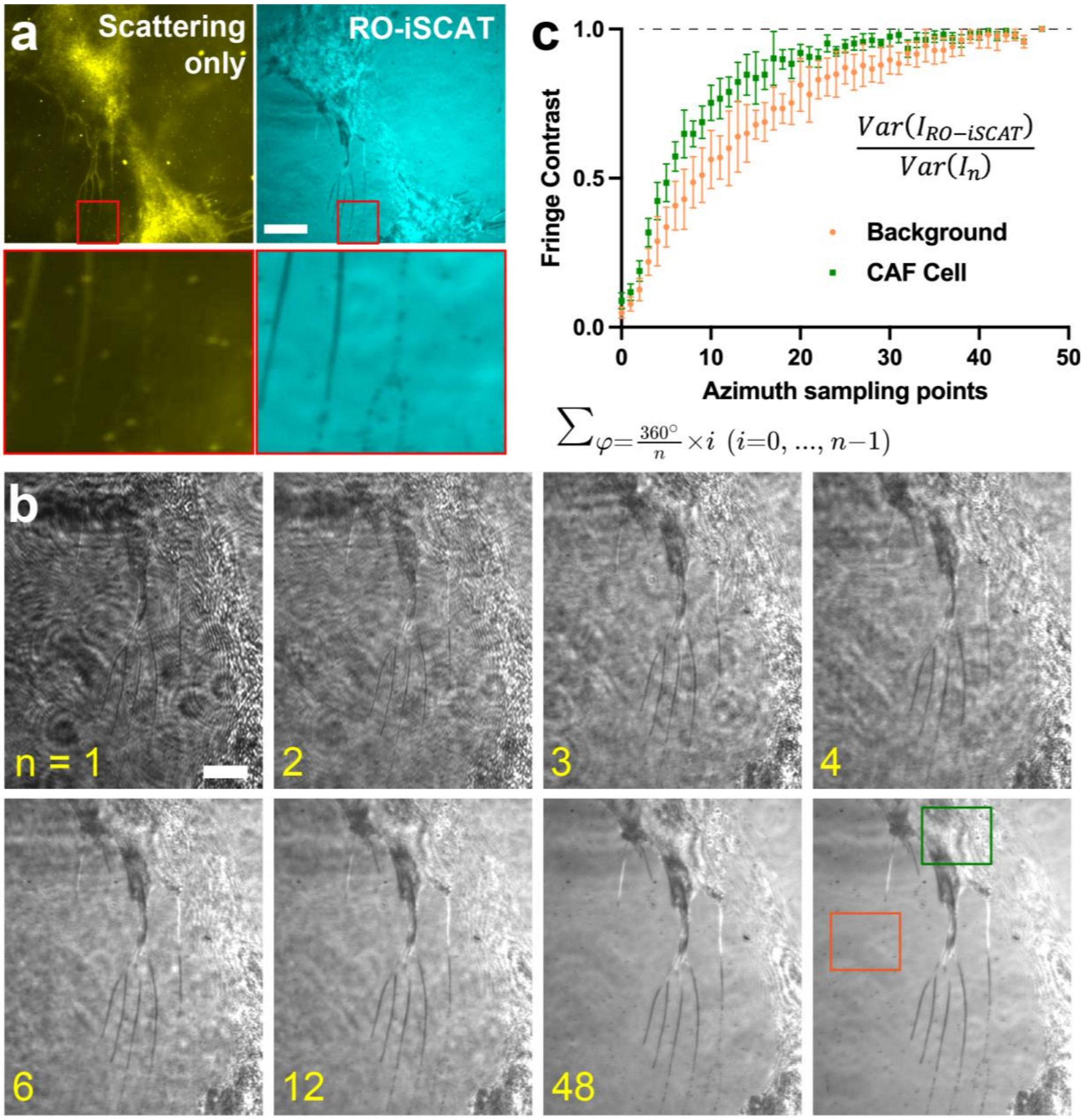


**Supplementary Fig. 5 Reduction of speckle background through rotational integration**. **a)** Scattering vs. RO-iSCAT modality of CAF cells, focusing on the protrusion and filopodia. **b)** Interference fringes under varying azimuth sampling points. **c)** Quantification of fringe contrast among regions of a CAF cell and background chosen from **b)**, e.g. the red and green rectangles in **b)**. **Scale bars**: **a)** 5 µm, **b)** 3 µm.

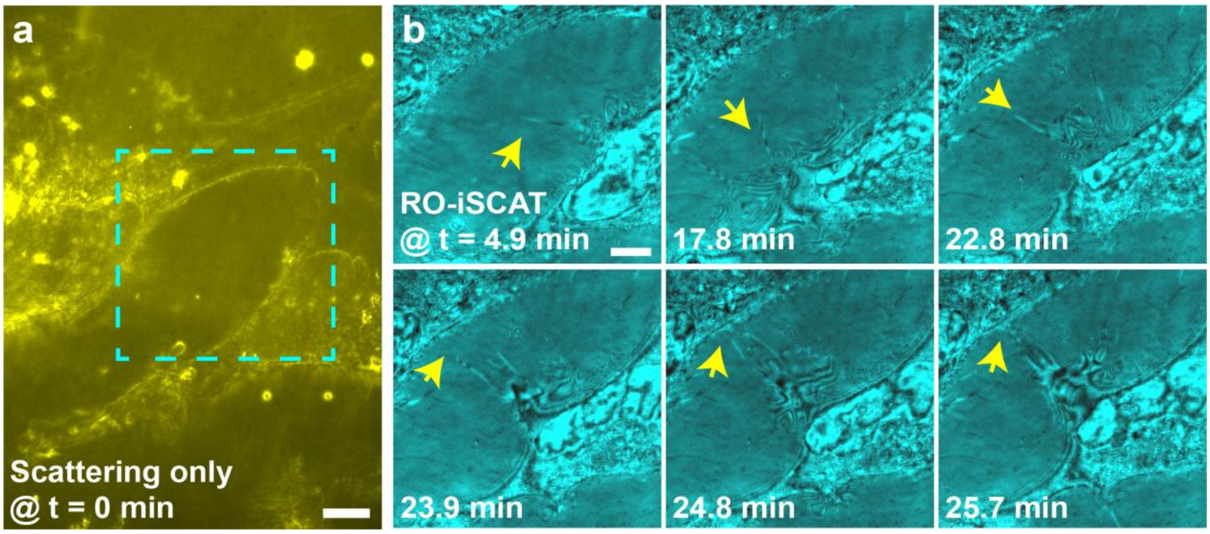


**Supplementary Fig. 6 Filopodium growth and external connections in endothelial cells**. **a)** A scattering snapshot at the initial time-point. **b)** Time-lapse RO-iSCAT at later time points corresponding to the cyan dashed box in **a)**. Main fringe variations are marked by yellow arrows. **Scale bars**: **a)** 3 µm, **b)** 2 µm.

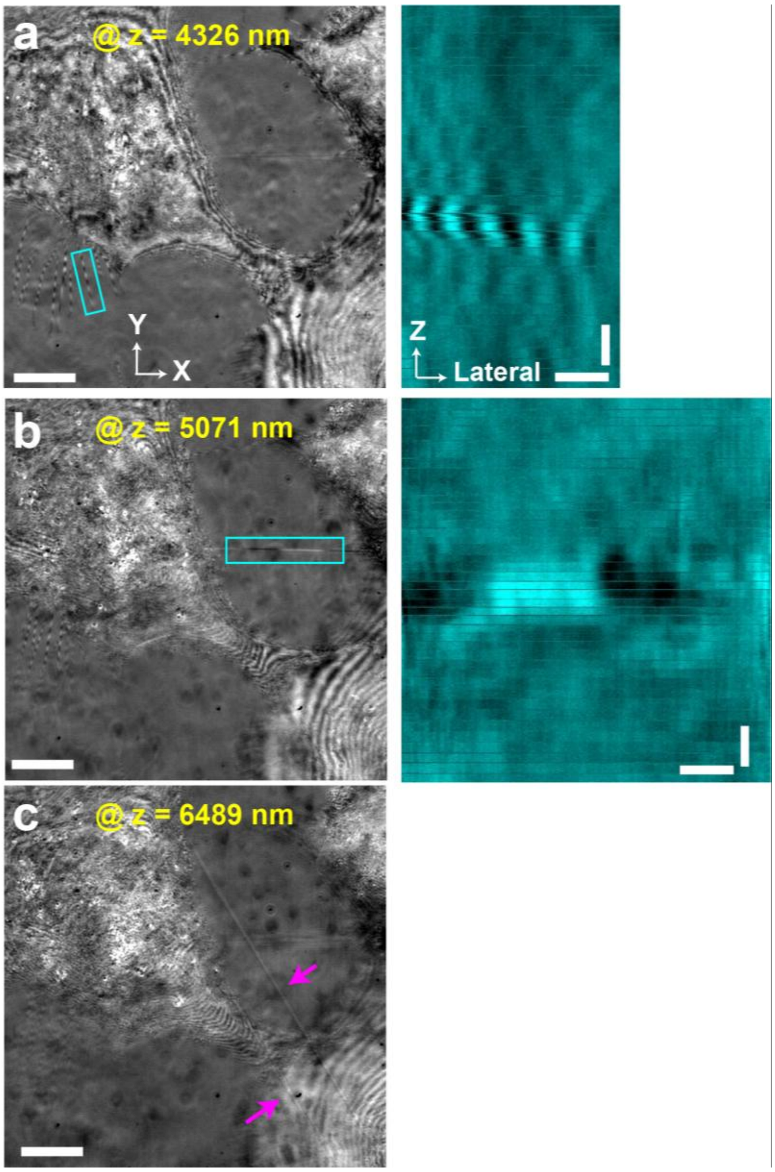


**Supplementary Fig. 7 Relative axial position of Type II and III membrane protrusion. a)** Resliced z-stack of cyan rectangle regions in **a)**, where the axes of slices are determined by the initial z axis and the long side of the rectangle. **b)** Resliced stack of a Type III protrusion. **c)** Long straight Type III membrane protrusion pointed out by purple arrows. Note that this z-sectioning stack was captured by axially moving the objective lens. When axially moving the objective lens, the interference pattern will not perform z-related intensity change, while both the axial movement of the sample dish and the inner axial difference will create intensity variance. **Scale bars**: **a)-c)** 3 µm, right insets in **a)**, **b)**, 1 µm.

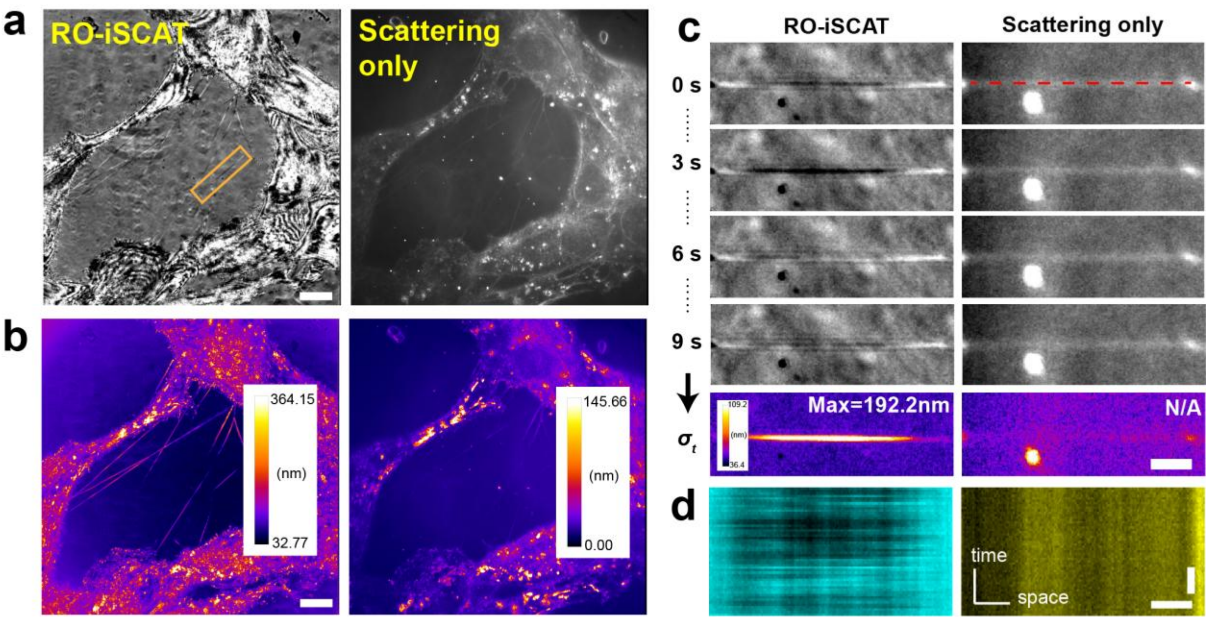


**Supplementary Fig. 8 Comparing intensity variation on membrane protrusion under RO-iSCAT versus scattering-only modality**. **a)** RO-iSCAT and scattering-only image on membrane bridges on CAF cells. **b)** Corresponding axial variation map of **a)**. **c)** RO-iSCAT versus scattering-only image at 0 s, 3 s, 6 s, 9 s time points and the generated intensity variation. The FOV corresponds to the orange box in **a)**. **d)** Kymograph along the red dashed line on membrane bridge in **c)**, separately from RO-iSCAT and scattering-only modality. **Scale bars**: **a)**, **b)** 3 μm, **c)** 2 μm, **d)** horizontal, 2 μm, vertical, 2 seconds.

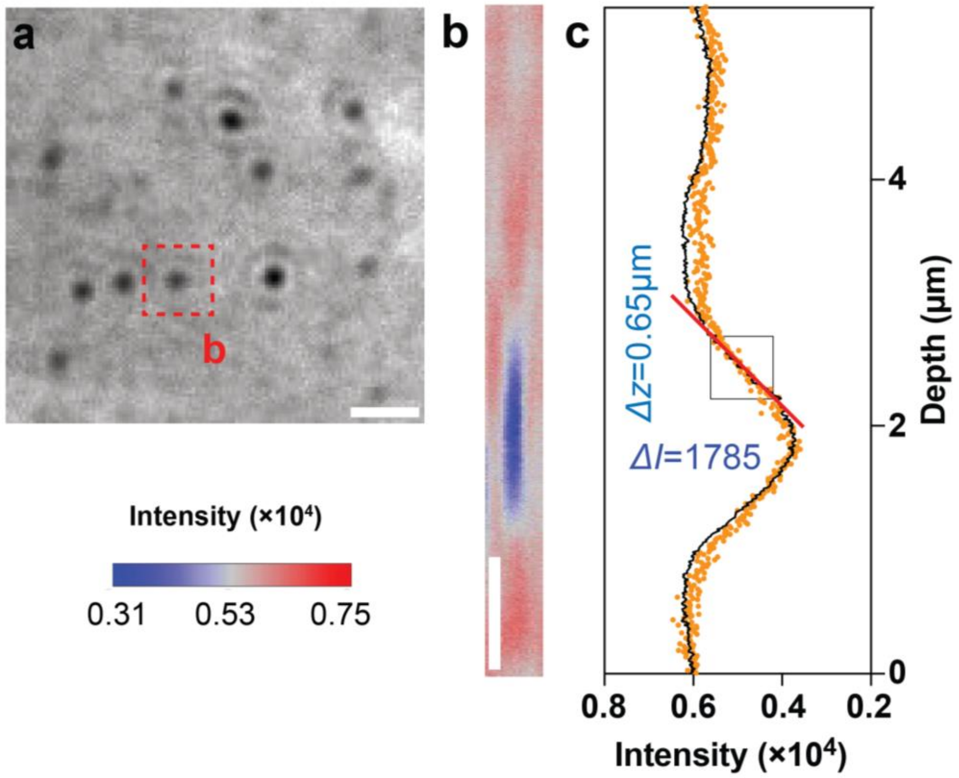


**Supplementary Fig. 9 Calibrating the intensity and axial displacement with 40nm nanoparticles**. **a)** RO-iSCAT image of 40nm particles in-focus. **b)** Axial section of the red dashed box in **a)**. **c)** Profile of intensity with depth. Orange scatters show the axial intensity along the particle in **b)** while the black curve is the averaged intensity profile among 8 particles in **a)**. The red line and black box indicate the linear region for calibration. **Scale bars**: **a)** 0.5 μm, **b)** 1 μm.

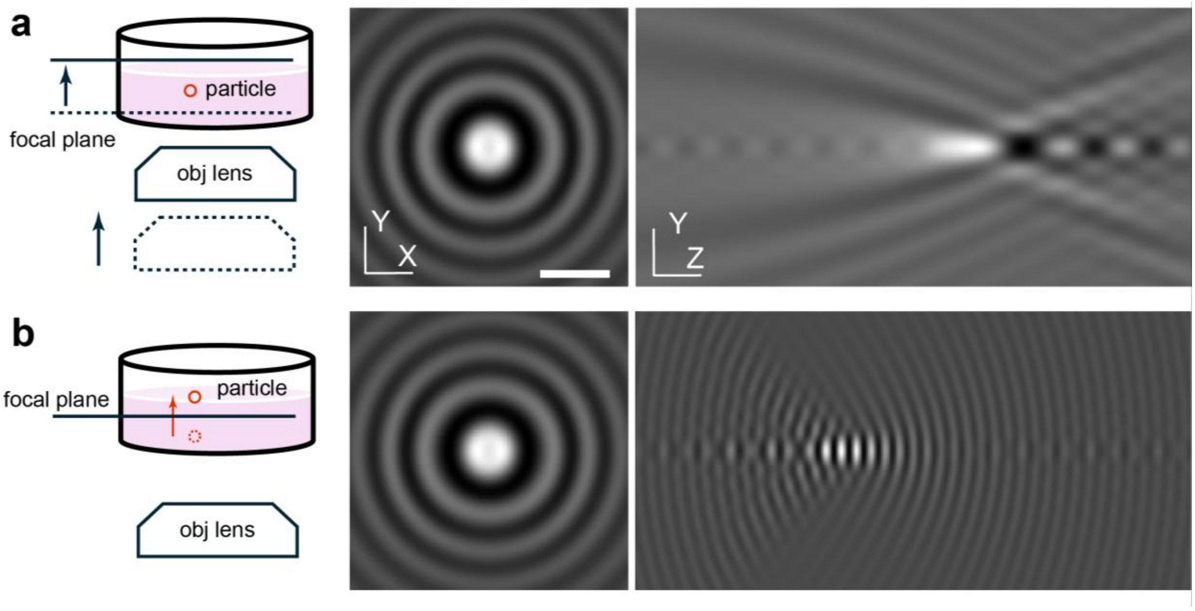


**Supplementary Fig. 10 iPSF comparison between two axial moving ways**. **a)** XY and YZ section of iPSF generated by the movement of focal plane, adjusting either the objective lens or the whole sample container. **b)** XY and YZ section of iPSF generated by moving single particle along principal optical axis. **Scale bars**: 1 μm.
